# Widespread 3’ UTR splicing regulates expression of oncogene transcripts in sequence-dependent and independent manners

**DOI:** 10.1101/2024.01.10.575007

**Authors:** Jack J. Riley, Cristina N. Alexandru-Crivac, Sam Bryce-Smith, Stuart A. Wilson, Ian M. Sudbery

## Abstract

**Background:** Splicing in 3’ untranslated regions (3’ UTRs) is generally considered a signal to elicit transcript degradation via nonsense-mediated decay (NMD) due to the presence of an exon junction complex (EJC) downstream of the stop codon. However, 3’ UTR intron (3UI)-containing transcripts are widespread and highly expressed in both normal tissues and cancers.

**Results:** Here we present and characterise a novel transcriptome assembly built from 7897 solid tumour and normal samples from The Cancer Genome Atlas. We identify thousands of 3UI-containing transcript isoforms, many of which are expressed across multiple cancer types. We find that the expression of core NMD component UPF1 negatively correlates with global 3UI splicing between normal samples, however this correlation is lost in colon cancer. We find that 3UIs found exclusively within 3’ UTRs (*bona-fide* 3UIs) are not predominantly NMD-sensitising, unlike introns present in 3’ UTRs due to premature termination. We identify HRAS as an example where 3UI splicing rescues the transcript from NMD. *Bona-fide*, but not premature termination codon (PTC) carrying 3UI-transcripts are spliced more in cancer samples compared to matched normals in the majority of cancer types analysed. In colon cancer, differentially spliced 3UI-containing transcripts are enriched in the canonical Wnt signalling pathway, with CTNNB1 being the most over-spliced in colon cancer compared to normal. We show that manipulating Wnt signalling can further regulate splicing of Wnt component transcript 3’ UTRs.

**Conclusions:** Our results indicate that 3’ UTR splicing is not a rare occurrence, especially in colon cancer, where 3’ UTR splicing regulates transcript expression in EJC-dependent and independent manners.

## Background

3’ untranslated regions (3’ UTRs) play essential roles in post-transcriptional gene regulation [1], including the regulation of transcript localization [2,3], stability [4,5], and translation efficiency [6,7]. Regulation of these processes can be attributed to the presence of cis elements within the 3’ UTR, for example: microRNA recognition elements (MREs) and RNA binding protein (RBP) binding motifs, which lead to the action of trans-regulators such as miRNAs and RBPs. Given the important regulatory roles of the 3’ UTR, it is unsurprising that cells regulate their 3’ UTR content in cell-type and condition specific manners, with classic examples being 3’ UTR lengthening during neuronal differentiation [8–10] and shortening in cancers [11–13]. The content of the 3’ UTRs can be regulated by 2 main mechanisms: alternative polyadenylation (APA) and alternative splicing (AS). APA relies upon utilisation of either distal or proximal polyadenylation sites to shorten or lengthen 3’ UTRs. AS also regulates 3’ UTR content but is not limited to the terminal sequence, instead alternate terminal exons might be used, or alternative 5’ splice sites and inclusion of earlier cassette exons might lead to the use of earlier stop codons. In this study we are interested in whether internal content can be spliced out of 3’ UTRs, which could be particularly useful in certain biological contexts where differential post-transcriptional regulation is required to impart additional functionality.

However, 3’ UTR splicing is generally seen as a signal to elicit nonsense-mediated decay (NMD), due to the presence of an exon junction complex (EJC) downstream of the stop codon [14,15]. EJC-linked NMD has previously been studied in yeast [16], humans [17,18], and zebrafish [19]. Thus, transcripts with spliced 3’ UTRs have generally been considered transcriptional noise and assumed to be broadly non-functional. Indeed, the GENCODE project labels transcripts with stop codons > 50nt upstream of a splice site as non-protein coding without further evidence, unless they are the only isoform of a known protein coding gene. However, there are a limited number of specific examples where eliciting NMD to produce short half-life transcripts may be functionally beneficial. For example, multiple SRSF (SRSF1-12) transcripts contain ultra-conserved poison exons which are subject to inclusion or excision via AS to modulate NMD and dampen expression, thereby maintaining steady state SRSF protein expression [20]. Additionally, a 3’ UTR intron (3UI) within the Arc mRNA leads to its degradation by NMD [21]. Whilst most Arc mRNA is processed and translated upon nuclear export, it has been reported that a subset of Arc mRNA is transported to neuronal dendrites in a translationally silent state, where, upon BDNF signalling, it is rapidly translated and degraded shortly thereafter, with such rapid bursts of expression and degradation potentially contributing towards long term potentiation and memory consolidation [22].

Whilst other specific examples of 3’ UTR splicing mediated regulation exist [23], they are limited to static biological contexts and do not address whether 3’ UTR splicing changes in different systems. In this study we investigate how 3’ UTR splicing impacts post-transcriptional regulation in cancer and how this differs from normal tissue. We achieve this by analysing a cohort of 7897 primary tumours and normal samples from The Cancer Genome Atlas (TCGA) as well as samples from the Cancer Cell Line Encyclopedia. Through the creation of 3UI-centric transcript assembly we investigate whether 3’ UTR splicing triggers NMD in the same way a premature termination codon does. Additionally, we investigate how 3’ UTR splicing regulates the composition of cis elements and the binding of trans-factors, and the effects of these changes on mRNA export and stability.

## Results

### A large set of highly expressed 3’ UTR intron (3UI) containing transcripts identified from cancer and normal RNA-seq data

Examining transcript annotations from Ensembl, we identified 8,566 transcripts with introns after the stop codon (4.5% of the Ensembl annotation). We noticed that many of these transcripts were almost identical to a matching isoform, however appeared to contain an early termination codon meaning that introns that would normally be in the coding region were now found after the stop codon and were annotated as within the 3’ UTR. To identify transcripts with *bona-fide* 3UIs (those that only ever occur in the 3’ UTR and never overlap the coding region), we filtered out transcripts where a putative 3UI had the same splice donor or splice acceptor location as an intron in the coding sequence of another transcript. A schematic overview of our transcript classification is shown in Figure 1A.

**Figure 1.**
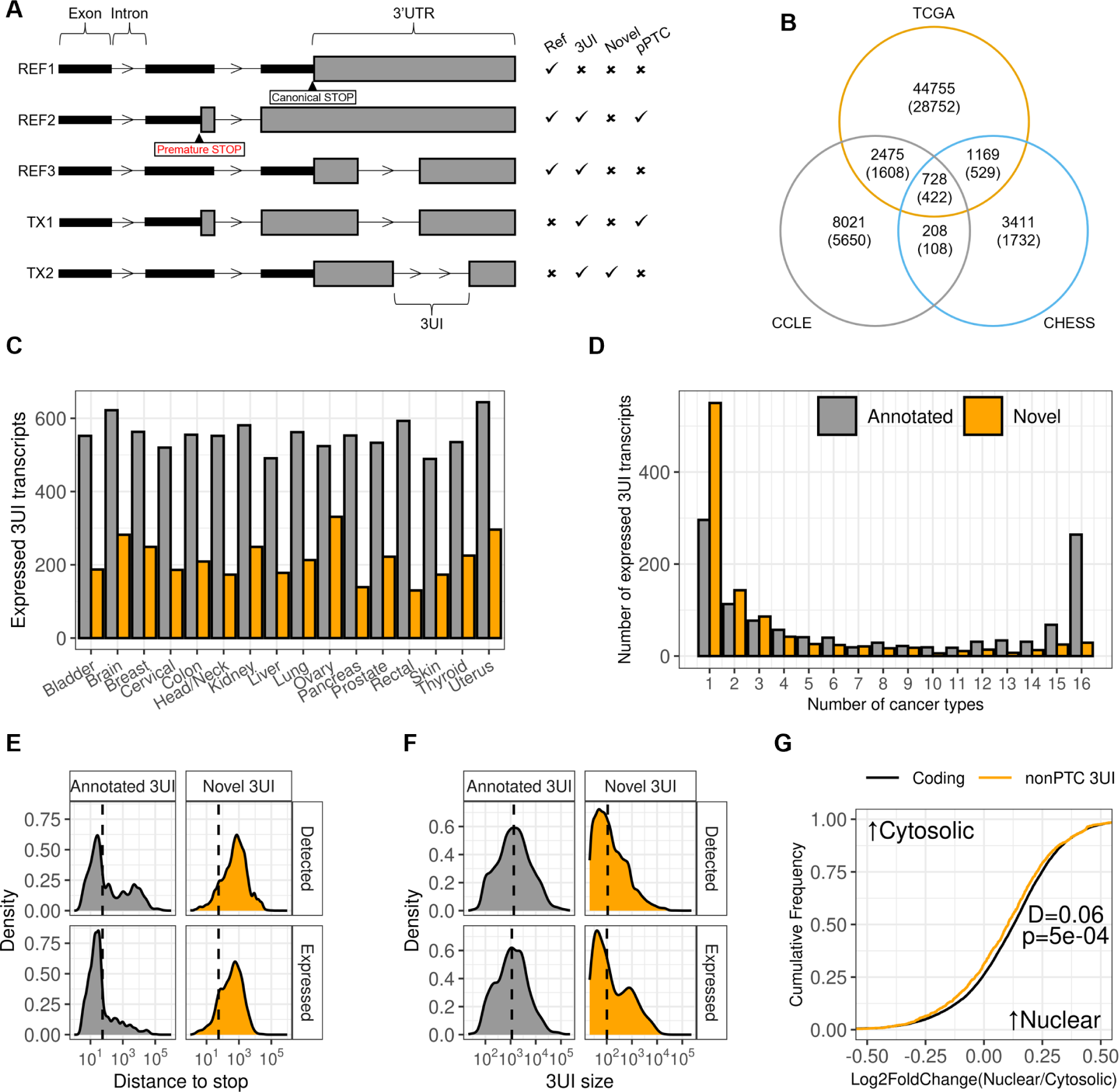
Detection and characterization of 3UI-containing transcripts. (A) Schematic diagram depicting the classifications of 3UI-containing transcripts. REF1 is a reference transcript lacking a 3UI, REF2 is a pPTC reference transcript, REF3 is a 3UI-containing reference transcript, TX1 is a novel-3UI containing transcript which has a CDS exon- and intron-chain identical to that of REF1, TX2 is a novel-3UI containing pPTC transcript which has a CDS exon- and intron-chain identical to that of REF2. (B) Comparison between the total number of 3UI-containing transcripts detected in the TCGA assembly, CCLE assembly and CHESS (GTEx) assembly, numbers in brackets refer to nonPTC transcripts. (C) Bar plot displaying the frequency that 3UI-containing transcripts are expressed in multiple cancer types. Transcripts are considered expressed where TPM > 1 and transcript/gene fraction > 0.25 in at least 10% of normal or cancer samples (see Materials and Methods). Grey bars represent transcripts containing annotated 3UIs. Orange bars represent transcripts containing novel 3UIs. (D) Bar plot comparing the number of expressed 3UI-containing transcripts between different tissues. Grey bars represent transcripts containing annotated 3UIs. Orange bars represent transcripts containing novel 3UIs. (E) Density plots comparing the distance between the start of the 3UI and stop codon (or maximal distance where there are multiple 3UIs) between annotated and novel 3UIs in expressed transcripts vs total detected. (F) Density plots comparing the size of 3UIs between annotated and novel 3UIs in expressed transcripts vs total detected. (G) Empirical cumulative distribution function plot comparing nuclear-cytosolic distribution of nonPTC 3UIs compared to protein-coding transcripts in HCT116. Only transcripts expressed TPM > 1 were considered. Left-shift represents more cytosolic localization, right-shift represents more nuclear localization. Kolmogorov-Smirnov test was performed to produce D and P statistics, P<0.05 is considered statistically significant.

To gain a more expansive view of splicing in 3’ UTRs, we examined the CHESS transcript annotation which is derived from normal tissue samples from the GTEx dataset. As these transcripts do not have an annotated coding sequence (CDS), we compared them to a high confidence set of Ensembl transcripts (see materials and methods). We identified transcripts that contained the full and unaltered CDS of an Ensembl transcript. Any introns in these transcripts after the stop codon we called 3UIs. If no 3UI had splice donors or acceptors that are used by any coding transcript, we marked the transcript as “non-PTC 3UI containing”. Finally, if the 3UI is found in no transcript in the reference annotation we labelled it a “Novel 3UI” (Figure 1A). We identified 5,516 3UI-containing transcripts, 2,791 of which were non-PTC and 948 novel in the CHESS set (Figure 1B, Supplementary Data 1,2).

It has been well established that cancerous cells show dysregulated patterns of splicing. In order to identify more examples, we assembled transcripts from 7,897 tissue samples from 16 solid tumour types in TCGA, and 348 samples from the Cancer Cell Line encyclopaedia (CCLE). After filtering (see materials and methods) we identified a set of 49,127 transcripts from TCGA and 11,432 transcripts from CCLE carrying 3UIs, 31,311 and 7,788 of which respectively were non-PTC (Figure 1B, Supplementary Data 3-6). Notably, these 3UIs utilised canonical donor-acceptor (GT-AG) splice sites 84.1% of the time (Supplementary Figure 1A) and had splice-sites that showed increased conservation compared to surrounding sequence (Supplementary Figure 1B). We studied the completeness of our experiment by examining subsets of colon cancer samples, showing that in this cancer type, the total number of 3UI containing transcripts identified began to saturate at 100 samples (Supplementary Figure 2A). Since our combined TCGA assembly contains over 78x this number, we expect that we are approaching the limit of 3UI containing transcripts that can be identified this way.

To guard against the possibility of generating spurious 3UI containing transcripts, we simulated 60 RNA-seq experiments using the Ensembl reference and passed these through our pipeline. We found that a small number of novel 3UI containing transcripts were assembled. However, the number of transcripts found began to saturate by 20 samples, suggesting we had identified the majority of these artifactual 3UI transcripts (Supplementary Figure 2B), which were subsequently excluded from further analysis.

We defined a threshold for transcripts being expressed as having a transcript expression level of at least 1 TPM and requiring that the transcript represented at least 25% of all transcript expression from a gene (25% Tx/G). Across all samples 6,934 non-PTC transcripts were expressed at this level in at least one sample. Applying these thresholds to samples from colon cancer, we found that many transcripts identified are expressed in only a single sample (Supplementary Figure 3). However, 764 transcripts reached this level in at least 10% of normal or colon cancer samples, these transcripts we describe as “broadly expressed” (Supplementary Figure 3, Supplementary Data 8). Despite the overall larger number of novel transcripts, more transcripts with previously annotated introns were classed as broadly expressed, and this pattern held across all cancer types examined (Figure 1C). Broadly expressed transcripts with novel introns were mostly only associated with one cancer type. By contrast, transcripts carrying previously annotated introns were broadly expressed in either a tissue-specific manner, or in all cancer types examined (Figure 1D).

We benchmarked our assembly using long-read RNA-seq data from the ENCODE project. Overall, 53% of the non-PTC 3UIs from broadly expressed transcripts in our TCGA dataset overlapped with non-PTC 3UIs in the ENCODE long read data. As the ENCODE long read data was derived from 264 cell lines, rather than primary cancer patient samples, we compared ENCODE long-read data for the colon carcinoma cell line HCT116 with 3UIs from transcripts in our dataset that were expressed at more than 1 TPM in HCT116 cells. In total 73% of such 3UIs overlapped with 3UIs in the ENCODE set.

It is generally thought that transcripts with splice junctions more than 55nt from the stop codon are sensitive to NMD. For previously annotated non-PTC junctions, most junctions were closer than 55nt from the annotated stop codon. This was not the case for the novel junctions. These were on average substantially further from the stop codon than 55nt, which would be expected to make them sensitive to NMD (Figure 1E, top). Surprisingly, this was not substantially different for transcripts broadly expressed in colon cancer (Figure 1E, bottom). This suggests that either these transcripts have some mechanism for avoiding NMD, or that these transcripts are so highly expressed that a large amount of transcript is present despite being subject to decay. Novel introns were also substantially shorter than annotated ones, including those which are broadly expressed in colon cancer (Figure 1F). Finally, to determine whether these transcripts are exported from the nucleus normally, we compared expression in the cytoplasm and nucleoplasm of HCT116 cells. We found that not only are non-PTC 3UI containing transcripts efficiently exported from the nucleus, but their nuclear:cytoplasmic ratios are slightly, but significantly, lower than for non-3UI containing protein coding transcripts (Figure 1G).

### 3’ UTR splicing regulates nonsense-mediated decay in unexpected ways

Whilst it is widely accepted that NMD acts as an RNA surveillance mechanism to reduce the expression of truncated mRNA species, its role in regulating *bona-fide* (non-PTC) 3UIs is less well defined. Given that UPF1 is necessary for NMD, we predicted that its expression levels would negatively correlate with the proportion of spliced reads across 3’ UTR introns.

For each sample in the TCGA colon dataset we calculated an “average 3UI percent spliced out (PSO)” value per sample (see materials and methods). We then correlated this with the expression of each gene in each sample (see materials and methods) to produce a correlation coefficient for each gene.

As expected, in normal samples we observed a negative correlation between UPF1 expression and average 3UI PSO for both non-PTC 3UIs (Figure 2A) and 3UIs that overlap protein coding introns in other transcripts (pPTC 3UIs; Supplementary Figure 4B). Surprisingly, we observed a loss of this correlation in cancer samples (Figure 2A; Supplementary Figure 4B). Such a difference between normal and cancer samples was also observed for other NMD components including UPF3B, SMG1, SMG5, SMG6 and SMG7 (Supplementary Figures 4A,C). Whilst SMG8 expression displayed no significant correlation with average 3UI PSO in normal samples, it displayed a positive correlation with the average non-PTC 3UI PSO of the cancer samples. This suggests that NMD-mediated regulation of 3’ UTR spliced transcripts may differ between normal and cancer samples.

**Figure 2.**
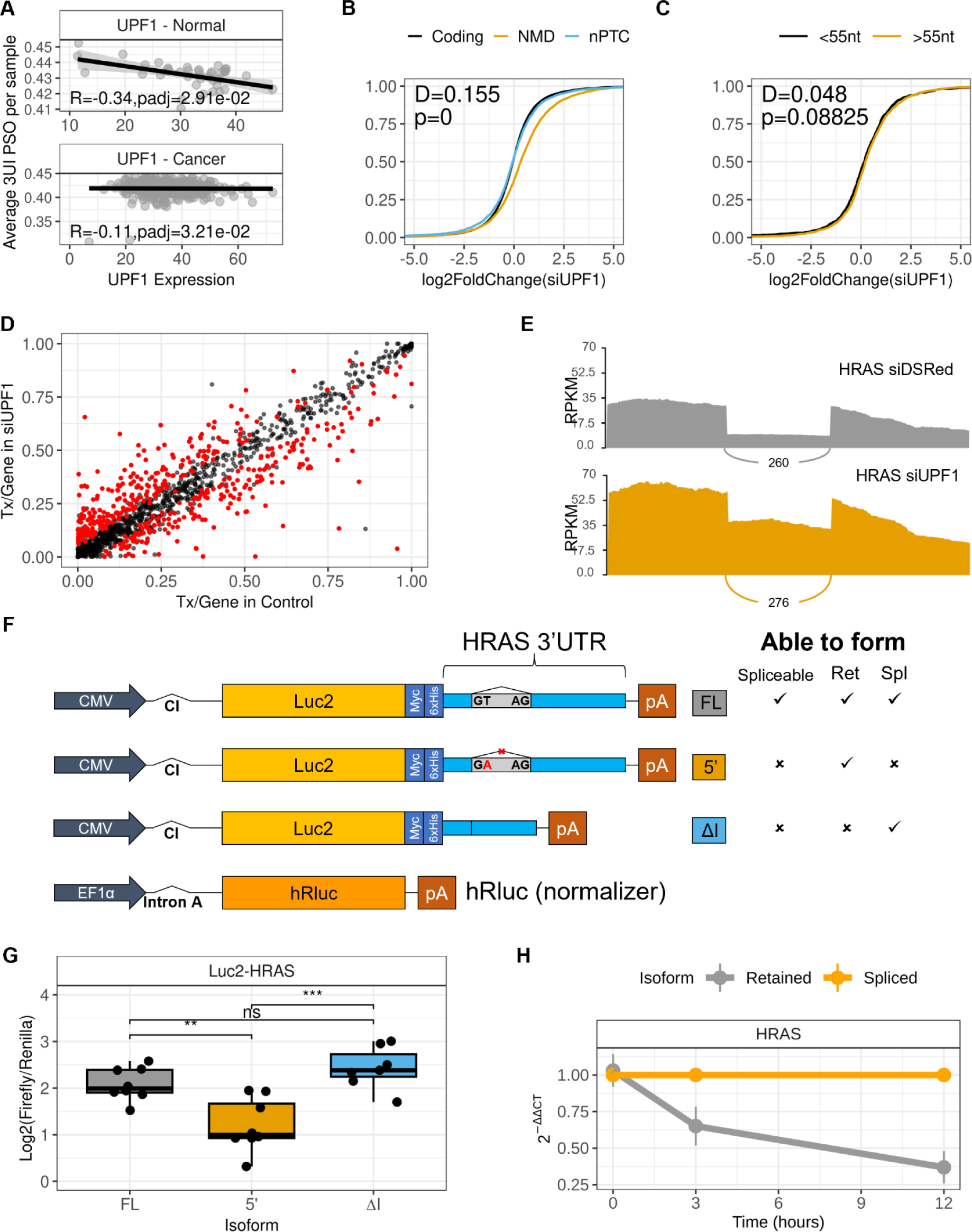
3UI splicing and retention regulates transcript sensitivity to nonsense-mediated decay. (A) Correlation of UPF1 expression in each sample with its corresponding average 3UI PSO. Strength of correlation was determined by calculating Spearman’s rank correlation coefficient. (B-C) Empirical cumulative distribution function plots comparing the expression changes induced by UPF1 knockdown on: (B) transcripts that are protein coding (black line), nonPTC (blue line) or expected to be NMD sensitive (pPTC transcripts and transcripts annotated as NMD-sensitive; orange line); (C) transcripts that have their terminal 3UI more (orange line) or less (black line) than 55 nucleotides from the termination codon; Kolmogorov-Smirnov test was performed, P<0.05 is considered statistically significant. (D) Differential transcript usage of 3UIs upon UPF1 knockdown. Each dot represents a 3UI-containing transcript. Red dots represent significant differential transcript usage (P<0.05, effect-size > 5%). (E) Sashimi plot comparing retained vs spliced read coverage for HRAS in siUPF1 (orange) and negative control siDSRed (grey). (F) Schematic diagram of Luciferase2-HRAS 3’ UTR reporter plasmids. Ticks and crosses indicate whether each construct is able to be spliced, and which isoforms it produces upon transfection. (G) Relative luminescence comparison between constructs upon transfection into HCT116 cells. (H) Relative stability of HRAS spliced and retained isoforms in HCT116 cells treated with Actinomycin D.

To investigate this further we performed RNAi against UPF1 in HCT116 cells (Supplementary Figure 5A-B) followed by RNA sequencing. UPF1 knockdown led to an increase in expression of traditional NMD targets, including transcripts labelled “Nonsense mediated decay” by Ensembl, and pPTC 3UI-containing transcripts from our assembly. However, this was not the case for transcripts with non-PTC 3UIs (Figure 2B). Given that an intron has to be further than 55nt from a stop codon to elicit NMD, we tested whether this result could be explained by the distance between stop codons and 5’ splice-sites. Surprisingly, we observed no significant difference between non-PTC 3UI carrying transcripts where the intron was more or less than 55nt from the stop codon (Figure 2C). These results confirm that 3UI-containing transcripts which are likely to contain a PTC are indeed NMD sensitive, however, *bona-fide* 3UIs may be NMD sensitive or insensitive (Figure 2D) irrespective of their position relative to 55nt after the stop codon.

Indeed, we find examples of splicing events that appear to rescue transcripts from NMD, such as in the oncogene HRAS (Figure 2E, Supplementary Figure 5C). Through conducting both differential splicing analysis (Figure 2E) and differential transcript expression analysis (Supplementary Figure 5D) we reveal a significant increase in the expression of the 3UI retaining isoform in siUPF1 compared to a control siRNA, whilst no change is observed in the 3UI spliced isoform. This suggests that UPF1 negatively regulates expression of the 3UI retaining isoform in colon cancer under non-knockdown conditions. This also suggests that splicing out the 3UI can offer protection from UPF1-mediated decay. To determine whether this effect is dependent on splicing, and therefore EJC deposition, we produced a set of Luciferase reporter constructs (Figure 2F) which contained either: the full-length HRAS 3’ UTR (Luc2-HRAS-FL) where the 3UI can be spliced or retained; the HRAS 3’ UTR with a GT>GA 5’ splice site mutation (Luc2-HRAS-5’) where the 3UI is always retained; or the HRAS 3’ UTR with the 3UI cloned out (Luc2-HRAS-ΔI). We confirmed that the correct isoforms are produced upon transfection of each plasmid into HCT116 cells (Supplementary Figure 6A). We observe that upon forcing 3UI retention there is a significant decrease in reporter activity, whilst removing the intron (via endogenous splicing of Luc2-HRAS-FL, or via cloning it out in Luc2-HRAS-ΔI) produces significantly more reporter activity (Figure 2G). Given that this significant increase in reporter activity is observed in the Luc2-HRAS-ΔI construct which is unable to be spliced by the cell and therefore does not have an EJC upstream of the intron, we conclude that this phenomenon is EJC-independent. Further, measuring mRNA stability following transcription inhibition with Actinomycin D reveals that the endogenous spliced isoform is more stable than its retaining partner (Figure 2H). This suggests that the splicing event is removing destabilising sequence elements present within the intron.

### 3’ UTR splicing is dysregulated in cancer

In order to investigate whether usage of 3UIs changes between normal and cancer samples, we used two complementary approaches: Differential exon usage (DEU) and differential transcript usage (DTU). In this case, differential exon usage measures the difference in usage of the retained intron that can be spliced out of the 3’ UTR. It has the advantage that it deals directly with the splicing event itself and is therefore closer to the mechanism. The disadvantages are: 1) It does not account for the rest of the transcript – e.g. does this splicing event change the total fraction of transcripts that have spliced 3’ UTRs or is splicing occurring in a transcript that already contains other 3UIs; 2) An intron can be significantly changed in usage, but the transcript it is being spliced from might only represent a minor part of the output from the locus in question (e.g. splicing in a 5TPM transcript of a 100TPM gene). DTU solves these problems, but has its own drawbacks: 1) the same 3UI may be present in multiple isoforms, thus a change in usage of one 3UI containing isoform does not necessarily imply a change in the total 3UI usage; 2) it is well known that global splicing is dysregulated in cancer, an increase in the number of splice isoforms used may lead to a decrease in the proportion of expression coming from any one isoform even if expression of the 3UI containing isoform itself is unchanged. Thus we used both approaches to study cancer related changes in 3UI usage.

Firstly by concentrating on the distribution of all changes (regardless of individual significance levels), across all 3UIs, most cancer types show a reduction of usage of the spliced isoforms in cancer for the majority of cases (Figure 3A, Supplementary Figure 7). This is consistent with a known increase in intron retention in cancer [24]. However, if we restrict our analysis to non-PTC 3UIs we see an increase in usage of the spliced isoforms in cancer for more transcripts than we see a reduction in the majority of cancer types (Figure 3A, Supplementary Figure 7). This suggests that *bona-fide* 3UIs behave in a qualitatively different way to introns that find themselves in the 3’ UTR by virtue of the inclusion of an upstream PTC, and that there is an increase in usage of *bona-fide* 3UIs in cancer.

**Figure 3.**
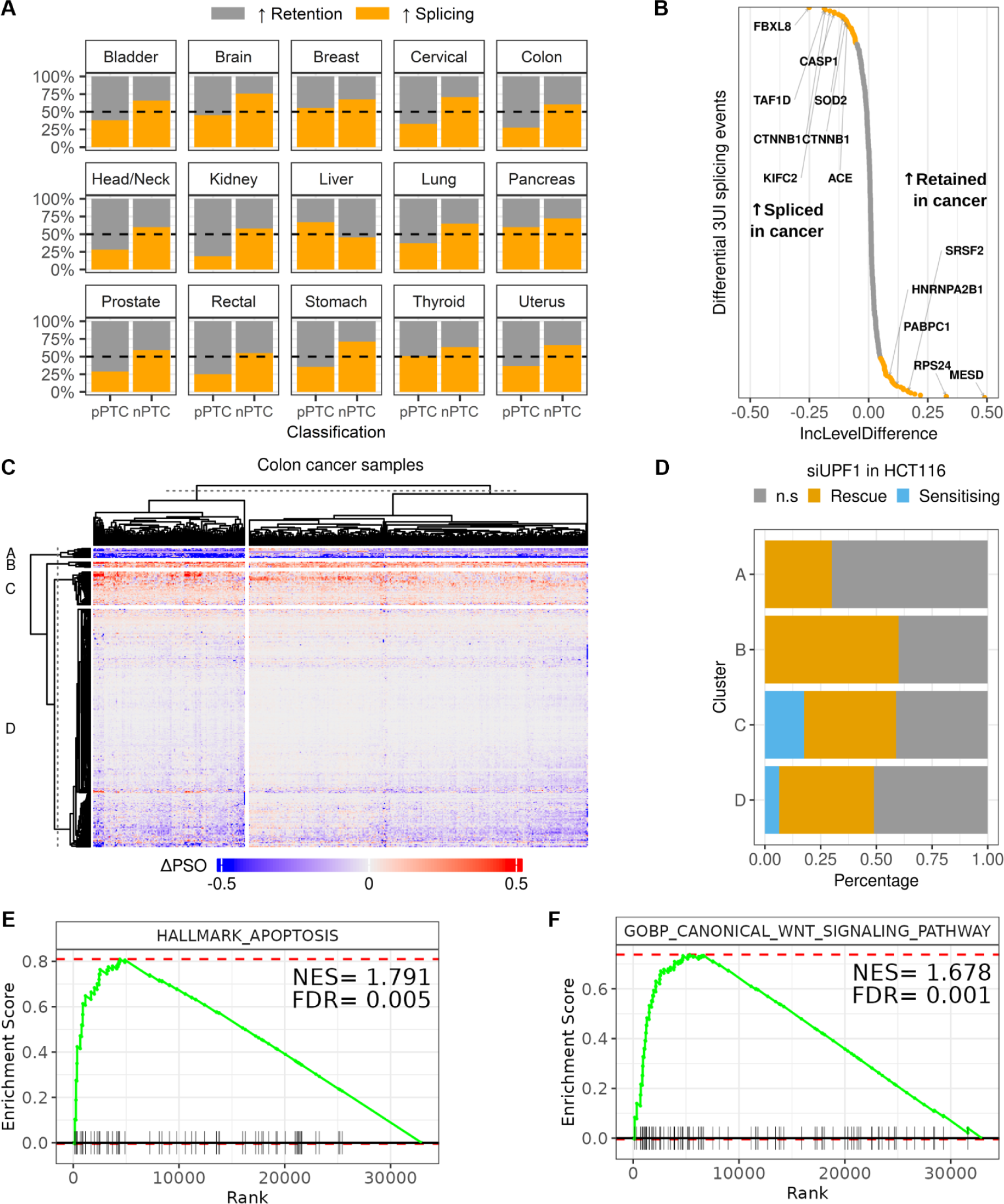
Widespread dysregulation of 3UI splicing in cancer. (A) Comparison of splicing of pPTC and nonPTC 3UIs between normal and cancer samples in a variety of tissue types from TCGA. Orange bars represent the percentage of transcripts that are spliced more in cancer compared to normal. Grey bars represent the percentage of transcripts that are retained more in cancer compared to normal. (B) Distribution of significant intron inclusion events between normal and cancerous colon tissue. Grey dots represent significant events (padj < 0.05) with no effect-size threshold. Orange dots represent significant events with IncLevelDifference > 5%. Positive IncLevelDifference means more retention occurs in colon cancer, negative values mean more splicing occurs in colon cancer. (C) K-means clustering of PSO values for events that were significantly different between normal and cancer samples. (D) Percentage of events in each cluster that were upregulated (Rescue) or downregulated (Sensitising) upon UPF1 knockdown. (E-F) Gene set enrichment analysis for events which are differentially regulated between normal and colon cancer samples: (E) Apoptosis from the MSigDB Hallmarks Collection; (F) Canonical Wnt Signalling Pathway from MSigDB C5 Collection.

Turning to individual examples of significantly changed 3UI usage (FDR<0.05), we see 123 introns with a significantly decreased inclusion level (i.e. increased splicing) and 217 introns with an increased inclusion level (i.e. decreased splicing) in colon cancer (Figure 3B). Of these 340 introns, 82 were contained within transcripts that also show DTU between cancer and normal conditions. Genes containing a 3UI with highly increased splicing include two different introns in the 3’ UTR of the gene encoding the WNT pathway regulator β-catenin (CTNNB1) and the inflammasome regulator Caspase-1 (CASP1). Genes containing a 3UI with highly increased retention in cancer include the gene encoding the poly-A binding protein PABPC1 and the WNT pathway regulator MESD. To account for potential differences in 3’ UTR splicing profiles within the population of colon cancer samples we conducted k-means clustering on these significant events (Figure 3C). Clustering of samples revealed that a subset of the population had a greater degree of splicing/retention in either direction. Additionally, we did not observe any link between APC/CTNNB1 mutation status and splicing signatures. Clustering of events revealed 4 distinct clusters: A) Highly retained more in cancer; B-C) Spliced more in cancer; D) Slightly retained more in cancer. We predicted that events which are highly retained more in cancer would be NMD sensitising, with intron retention acting as a potential mechanism to evade NMD. Likewise, we predicted that events which are significantly over spliced may be NMD rescuing. To test this we cross-referenced these events against our UPF1 knockdown in HCT116 (Figure 2) and noted that whilst clusters B and C do show an increased level of NMD rescue (i.e. increase in number of spliced reads upon treatment with siUPF1), cluster A does not show any selection of NMD sensitising events, whilst cluster D only shows a small percentage (Figure 3D). Together these results indicate that for events which are significantly different between normal and cancer samples, the introns which are spliced more in cancer are likely to be less sensitive to NMD, whilst those which are retained more in cancer are likely to be more sensitive to NMD.

To identify potential functional implications of increased 3’ UTR splicing in colon cancer we ranked each splicing event from most over spliced (lowest IncLevelDifference) to most over retained (highest IncLevelDifference) and performed Gene Set Enrichment Analysis (GSEA). When utilising the MSigDB Hallmarks collection we observed significant enrichment of gene sets related to apoptosis (Figure 3E; NES=1.791), DNA repair (NES=1.735), IL2/STAT5 signalling (NES=1.692), E2F targets (NES=1.672) and Myc targets (NES=1.441). Given that CTNNB1 harbours one of the most over spliced 3’ UTR introns between normal and colon cancer, and its function as a central regulator of the Wnt signalling pathway, we tested for enrichment of the canonical Wnt signalling pathway using the C5 (ontology) gene set and observed a significant enrichment (Figure 3F; NES=1.678, FDR=0.001).

### 3’ UTR splicing modulates RNA-RBP and RNA-miRNA interactions

To investigate how 3’ UTR splicing may be regulating the inclusion or excision of binding sites for miRNAs and RBPs we used data from the ENCORI platform, which contains AGO-CLIP and CLIP-seq data, to determine whether non-PTC 3UI sequences were enriched for such elements. Upon examination of the full non-PTC 3UI set from our transcript assembly (n=74585) the most significantly enriched RBP-RNA interaction is with TARBP2 (Supplementary Figure 8A), a subunit of the RNA induced silencing complex (RISC) required for miRNA processing. Notably both HNRNPU and HNRNPUL1 binding are also significantly enriched in this analysis. Additionally, we see significant enrichment of various miRNA-RNA interactions, with miR-324-3p being most significant (Supplementary Figure 8B). To gain an insight into which RBP-RNA and miRNA-RNA interactions may be most relevant in the colon cancer setting we repeated this analysis on only the non-PTC 3UIs which were classed as “broadly expressed” in colon cancer and again observed enrichment of TARBP2 and HNRNPUL1 binding (Figure 4A). Notably the most enriched RBP interaction within this analysis was the pre-mRNA splicing factor PRPF8, whilst spliceosomal components SF3B4, and RBM22 were also highly enriched (Figure 4A). We also found that TARBP2, SF3B4 and RBM22, alongside m^6^A demethylase FTO and RNA helicase DHX9 binding was enriched in non-PTC 3UIs which show significantly increased splicing in colon cancer patients compared with normal colon controls (Figure 4B). In this comparison we also observed a significant under-representation of several miRNA, with the most significant being miR-450a-5p (Figure 4C)

**Figure 4.**
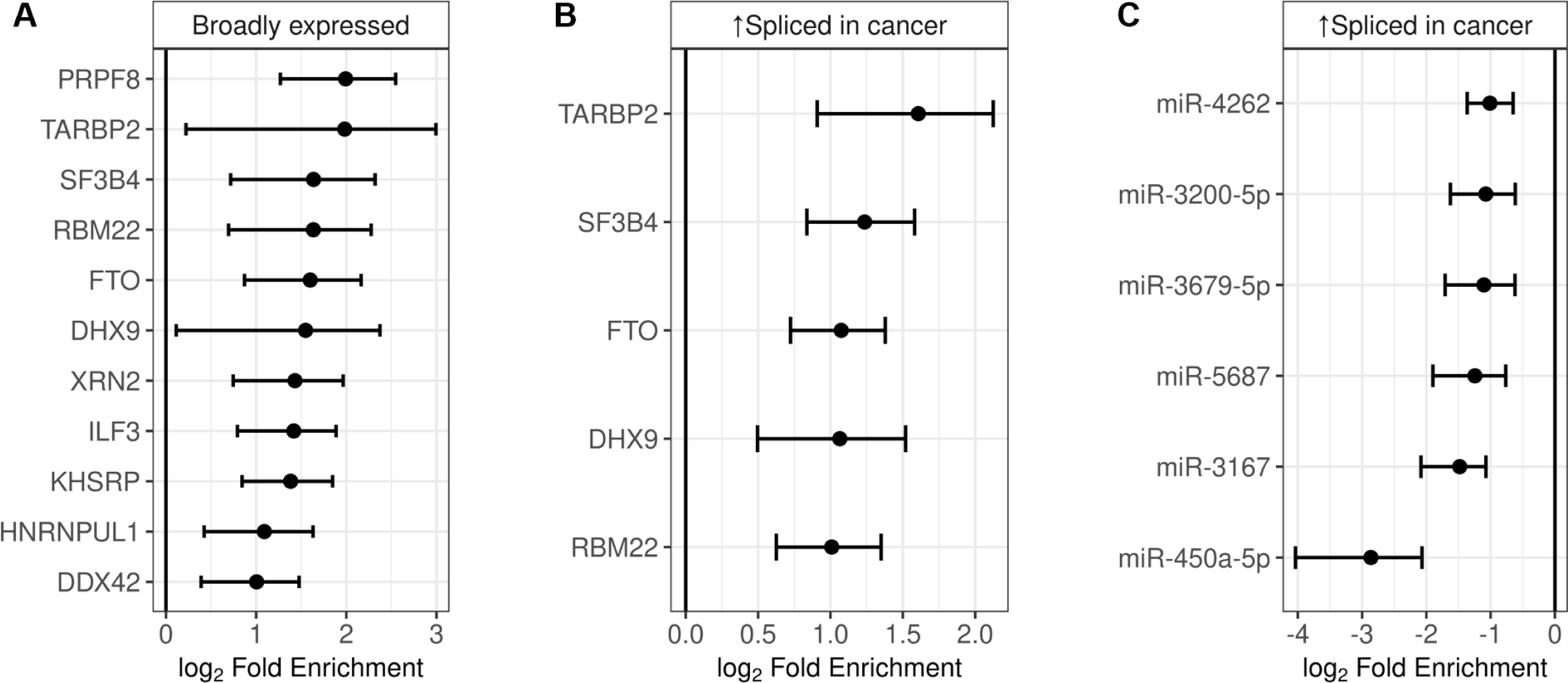
3UI splicing regulates RNA-RBP and RNA-miRNA interactions. (A-B) RBP enrichment analysis for 3’ UTR intron sequences. (A) 3’ UTR intron sequences from transcripts that are broadly expressed in colon cancer. (B) Sequences from 3’ UTR introns that are spliced more in colon cancer than normal samples. (C) miRNA under-enrichment in sequences from 3’ UTR introns that are spliced more in colon cancer than normal samples.

### Autoregulation of 3’ UTR splicing by the Wnt signalling pathway

Previous studies have shown that the Wnt signalling pathway is often dysregulated in colorectal cancer, in as many as 93% of samples [25], commonly leading to overactivation of CTNNB1, a central regulator of the canonical Wnt signalling pathway. CTNNB1 has two 3’ UTR-spliced isoforms, both utilise a common 5’ splice donor sequence but have alternate 3’ splice acceptor sequences. Thus the isoform utilising the proximal 3’ splice acceptor contains a 305nt long 3UI (short spliced isoform), whilst the isoform utilising the distal 3’ splice acceptor contains a 464nt long 3UI (long spliced isoform).

Further to our finding that 3UI splicing is dysregulated in cancers generally (Figure 3A) and specifically for CTNNB1 in colon cancer (Figure 3B), we postulated that this could be in part due to hyperactive Wnt signalling. To test this we treated HCT116 with varying concentrations of CHIR99021, a GSK3β inhibitor, to cause accumulation of nuclear β-catenin protein and therefore activate the canonical Wnt signalling pathway. Isoform-specific qPCR revealed dose-dependent regulation of the short spliced isoform, with no effect on the retained isoform (Figure 5A top) and observed the opposite with IWR-1, which stabilises Axin2 leading to β-catenin degradation (Figure 5A bottom). If this phenomenon was caused by alternative splicing, we would expect the expression of the retained isoform to displace the spliced isoform. However, this is not the case, therefore we suggest this is due to isoform-specific post-transcriptional regulation, e.g. regulation of mRNA stability.

**Figure 5.**
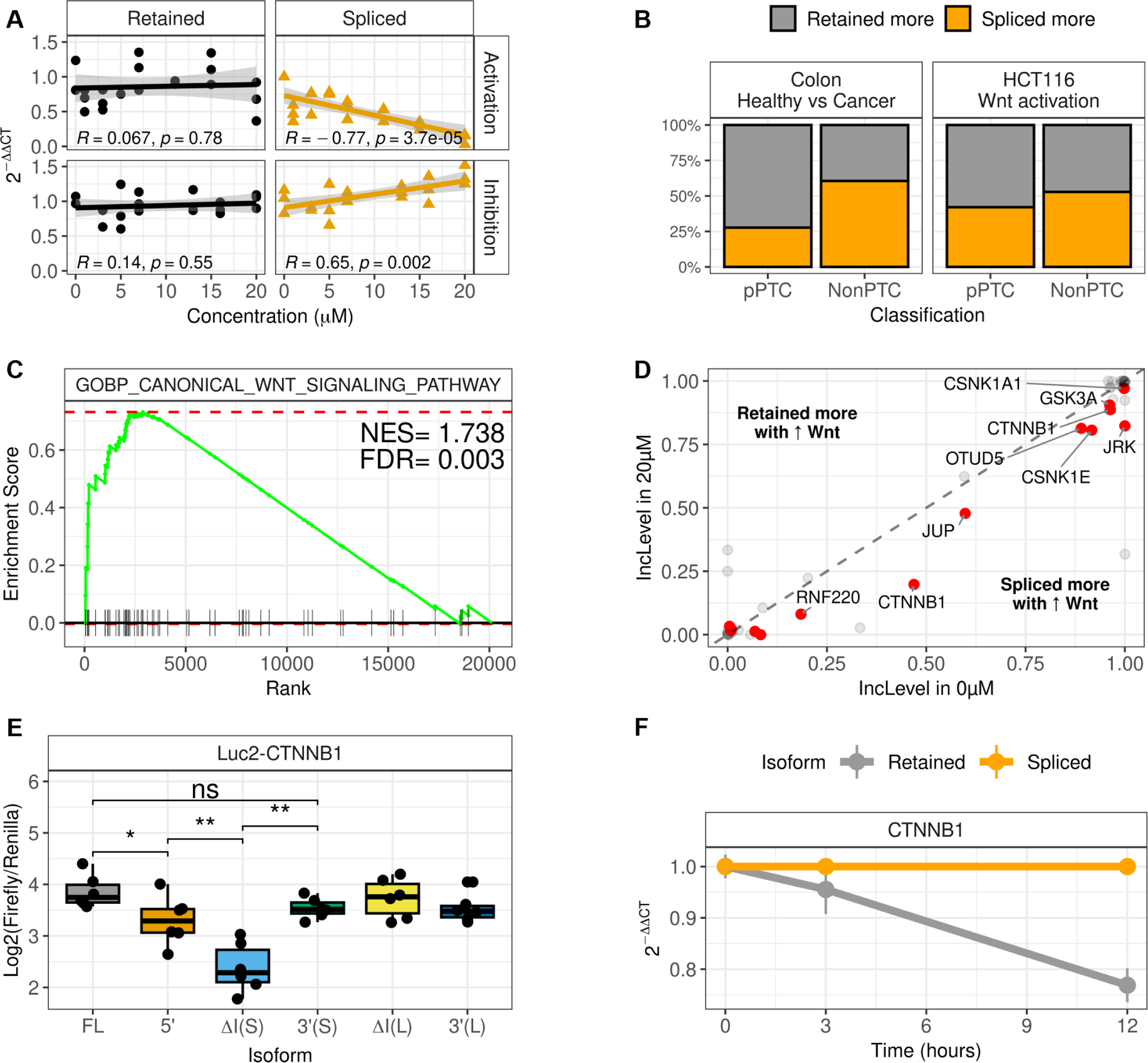
Wnt signalling regulates 3UI splicing of Wnt signalling pathway components. (A) Effect of increasing concentrations of CHIR99021 (Wnt signalling pathway activator) and IWR-1 (Wnt signalling pathway inhibitor) on expression of retained vs short-3UI spliced CTNNB1 isoforms in HCT116 cells. Pearson correlation coefficients and p-values are indicated. (B) Percentage of nonPTC and pPTC 3’ UTR introns that are spliced more (orange) or retained more (grey) upon Wnt signalling activation in HCT116 cells, compared with normal vs colon cancer samples. (C) Gene set enrichment analysis for events which are differentially regulated upon Wnt activation in HCT116 cells reveals enrichment of the canonical Wnt signalling pathway. (D) Differential intron inclusion of canonical Wnt pathway components in 0µM vs 20µM CHIR99021. Red dots represent significant events from RMATS (padj < 0.05). (E) Relative luminescence comparison between Luciferase2-CTNNB1 constructs (Supplementary Figure 6B) upon transfection into HCT116 cells. (H) Relative stability of CTNNB1 short-spliced and retained isoforms in HCT116 cells treated with Actinomycin D.

Next, we treated HCT116 with 20µM CHIR99021 and conducted RNA sequencing. Splicing analysis revealed that roughly equal proportions of 3UIs were spliced more or retained more, compared with the majority of 3UIs being over spliced in colon cancer (Figure 5B). Strikingly, we found that transcripts with increased 3UI splicing upon CHIR99021 treatment were strongly enriched in canonical Wnt signalling pathway components (NES=1.738, FDR=0.003, Figure 5C) as also observed between normal and colon cancer samples (Figure 3F). A breakdown of the Wnt signalling pathway components regulated by CHIR99021 treatment can be found in Supplementary Figure 9. Notably, the CTNNB1 long 3UI-spliced isoform, as well as a 3UI-spliced isoform of JUP (γ-catenin), were expressed significantly higher in CHIR99021 treated samples than untreated (Figure 5D). From these results we suggest that dysregulation of 3UI splicing in colon cancer may be in part due to hyperactive Wnt signalling.

Finally, to elucidate the difference between each CTNNB1 3’ UTR isoform we created Luciferase reporter plasmids as before (Supplementary Figure 6B-C). By mutating the 5’ splice site we observed a reduction in expression, however, when either of the 3’ splice sites are mutated this reduction is not observed (Figure 5E). Additionally, we show that when the splicing of one isoform is prevented (through 3’ss mutation) that splicing of the other isoform is enhanced (Supplementary Figure 6C), potentially explaining the lack of effect. Surprisingly, when the short intron is cloned out we observe a more drastic decrease in expression, yet when the long isoform is removed (which encompasses the short isoform) this result is not replicated. This can be explained by either: 1) removing the short isoform produces a binding site for a destabilising trans-factor, which cannot bind to the endogenously spliced isoform due to the presence of the EJC; 2) the short intron contains stabilising elements, but the long isoform contains additional destabilising elements that mask/counteract these. These are outlined in Supplementary Figure 10. Turning to the endogenous context in HCT116, we find that the 3UI-spliced isoform (short intron) is more stable than the 3UI-retaining isoform (Figure 5F). Together these results indicate that RNA stabilisation conferred by splicing the 3’ UTR of CTNNB1 is EJC-dependent, i.e. due to regulation of the mRNP.

## Discussion

Here we have presented a novel transcriptome assembly (Supplementary Data 7) containing thousands of 3UI-containing transcript isoforms ranging from patient-specific transcripts to transcripts that are broadly expressed in many or all cancer types analysed. This assembly has allowed us to characterise the extent of 3’ UTR splicing in both normal and cancer samples and has revealed that 3’ UTR splicing regulates transcript expression in both EJC-dependent and independent manners in colon cancer. We make both individual splice event and transcript level quantification data against this assembly publically available.

During the production of this manuscript a parallel analysis of 3’ UTR splicing in TCGA data was published [26]. Chan *et al* took a complementary approach to the transcript assembly approach taken here by identifying spliced reads that overlapped 3’ UTRs, but not coding sequences. Due to a lack of availability of the total dataset, we are unable to compare the transcripts we identified with the splice junctions they focused on. However, many of the findings are similar to ours at a global level. Our results complement and extend theirs in several key ways. Firstly, our approach allows a comparison of 3UIs that are always 3’ UTR located, with those that find themselves in a 3’ UTR due a premature stop codon. Secondly we examine the effect of knocking down key NMD regulator UPF1 on 3UI expression at a global level. Thirdly we show that not only do a large number of genes in the canonical WNT pathway appear to be regulated by 3UI splicing, but the activity of the pathway regulates 3UI splicing in turn.

Chan *et al* find that the majority of their events of interest are closer than 55nt to the stop codon, and suggest these events are therefore not NMD sensitive, demonstrating this for 5 example events. However, we find that many of our 3UIs are more than 55nt from the stop codon, suggesting that by conventional wisdom, they should be NMD sensitive.

It has previously been shown that cancers can exploit NMD to promote pro-oncogenic cellular behaviours [27,28] both through NMD-evasion by oncogenes [18] and NMD-sensitisation of tumour-suppressor genes [29]. It has also been shown that NMD efficiency can differ between cell types during cell differentiation [30], and even between individual cells within a population [31]. Here we provide evidence that UPF1-mediated regulation of transcripts differs between those containing PTCs compared with those which contain *bona-fide* 3UIs. Additionally, we show that for *bona-fide* 3UIs, the location relative to the threshold of 55nt past the stop codon has no significant effect on NMD sensitivity in colorectal carcinoma cell line HCT116. Given that EJCs would persist for 3UI events found >55nt from the stop codon, whilst they would be removed from the mRNP by the ribosome for 3UI events found <55nt, we suggest that any differences in UPF1 sensitivity between these 3UI containing transcripts are due to EJC-independent mechanisms.

UPF1 has also been shown to regulate the stability of transcripts with long 3’ UTRs through interaction with Ago2 and the miRNA pathway [33]. Indeed, we do find that some 3UIs serve to protect the transcript from decay. In the case of HRAS, the spliced isoform is more stable (Figure 2H) and the unspliced isoform, rather than the spliced isoform, is upregulated by UPF1 knockdown (Figure 2E, Supplementary Figure 5C,D). Splicing a 3UI could dramatically shorten the overall length of a 3’ UTR and rescue it from length-dependent decay. In HRAS the unspliced 3’ UTR is 260nt long, whilst the spliced 3’ UTR is 152nt long. This suggests that UTR-length mediated regulation may be more widespread than previously considered. miRNAs primarily regulate transcript stability through binding to the 3’ UTR, and 3UI retaining isoforms could harbour more MREs, therefore the increased expression of 3UI-retaining isoforms (such as in HRAS) upon UPF1 knockdown could be explained by UPF1-Ago2-mediated regulation.

Whatever the source of the effect, it appears that it is not EJC-dependent, as deletion of the HRAS 3UI, which prevents EJC-deposition while also removing the intron sequence, has little effect on expression of a Luciferase reporter, while mutation of the 5’ SS, which would also cause a loss of EJC-deposition, but does not remove the intron sequence, causes a reduction in expression (Figure 2G) compared to the full length, spliceable HRAS 3’ UTR. Whether this form of UPF1-Ago2-mediated regulation differs between non-cancer and cancer, or between different cancer types, remains unclear.

In contrast, the effects of splicing in the 3’ UTR of CTNNB1 do appear to be at least partially EJC-dependent, as mutation of the 5’ SS and deletion of the short intron both cause a reduction in expression (Figure 5E). These results differ from those of Chan *et al*, who find that both mutation of the 5’ SS and deletion of the long intron increase expression of a Luciferase reporter (we find that deletion of the long intron has no effect on expression)[26]. These differences could be explained by the different models used (Colorectal carcinoma here, and Hepatocellular carcinoma in Chan *et al*). Indeed, we find that the CTNNB1 generated by splicing the short intron is more highly used in Colorectal carcinoma samples than the isoform generated by splicing the long intron. However, our results do not provide support for the hypothesis that splicing in the 3’ UTR of CTNNB1 regulates expression by retention of the unspliced isoform in the nucleus via its interaction with U1, at least in HCT116 cells, as we would then not expect our 5’ ss mutant to reduce expression.

We propose a model for transcript stabilisation by 3’ UTR splicing in colorectal carcinoma both by EJC-independent and EJC-dependent mechanisms (Supplementary Figure 11). Regarding EJC-independent mechanisms, removal of 3UIs can remove destabilising sequence elements such as MREs and DRACH motifs which leads to less miRNA binding and less m^6^A modification (note the enrichment of m6a demethylase FTO in 3UIs). Additionally, removing 3’ UTR introns likely plays a role in manipulating RNA secondary structure, which may impact RNA stability and translation [7], although a widespread study of 3’ UTR splicing on predicted RNA structure has not been conducted to date. By contrast, EJC deposition may protect local sequence from miRNA and RBP binding by steric interference, and does not always elicit NMD in colorectal carcinoma cells, as we have shown here. Additionally, it has recently been shown by two separate studies that EJC deposition appears to protect upstream and downstream sequence (∼100nt) from m^6^A modification as part of a so-called “m^6^A exclusion zone” [34,35], this has been shown to be in part due to EJC core component EIF4A3 [34]. Hence splicing the 3’ UTR would not only remove potential m^6^A sites (via DRACH motif exclusion), but then further protect the surrounding sequence from modification, to increase RNA stability. Therefore we propose that, at least in the colorectal carcinoma setting, that splicing 3’ UTRs is able to modulate the composition of the mRNP to modulate post-transcriptional regulation beyond just regulating NMD-sensitivity.

We have also shown that manipulation of the Wnt signalling pathway regulates alternative 3UI splicing of canonical Wnt signalling pathway component transcripts. However, whether this is due to direct action of the Wnt signalling pathway (perhaps through kinase or phosphatase activity) or due to genes regulated by TCF/LEF transcription factors remains unclear. Nevertheless, understanding the molecular mechanisms downstream of Wnt hyperactivation in cancers remains critical to discovering novel therapeutic avenues. Wnt activating mutations in CTNNB1 or inactivating mutants in APC have been identified in 80% of colorectal carcinoma samples from the TCGA cohort studied here [25], but Wnt signalling is also commonly hyperactivated in many other cancers [36,37] such as hepatocellular carcinoma [38], pancreatic cancer [39] and esophageal squamous cell carcinoma [40,41]. Increased Wnt signalling activity has been linked with “stem-like” behaviours including increased cell proliferation [42], therefore tight control of its action is crucial for tissue homeostasis, which is lost in cancers. It is possible that aberrant splicing regulation of Wnt components contributes to this phenomenon. Additionally, non-canonical functions of the Wnt signalling pathway, for example in planar cell polarity [43], and also additional functions of β-catenin, should be considered in the cancer setting given their roles in cell adhesion, migration and invasion [44,45]. In this regard we also show that JUP (γ-catenin, also known as junction plakoglobin) 3’ UTR splicing is regulated by Wnt signalling. JUP has been shown to function in a compensatory manner with β-catenin at adherens junctions [46,47].

## Conclusions

We conclude that splicing in the 3’ UTR is not a rare occurrence, especially in colon cancer. 3’ UTR splicing events represent more than transcriptional noise, and their inclusion or exclusion in different cellular contexts via alternative splicing represents a mechanism to regulate transcript expression by both EJC-dependent and independent mechanisms.

## Methods

### Identification and quantification of 3UI expression

Raw RNAseq reads from The Cancer Genome Atlas Program (TCGA) [48] or the Cancer Cell Line Encylopedia (CCLE) [49] were downloaded in .fastq format. Read quality control was conducted with FastQC to ensure [50]. Reads aligned to hg38 for TCGA samples passing QC were obtained from the Genomic Data Commons (dbGaP accession phs000178.v11.p8) in BAM format. Raw reads for CCLE samples were mapped to hg38 with STAR [51] using junction annotations from ENSEMBL 85.

3UI detection was facilitated via a custom bioinformatic pipeline (www.github.com/sudlab/Cancer3UIs/pipelines/pipeline_utrons_assemble.py). Briefly, reads were first filtered by Portcullis [52] to retain only the spliced reads that are likely to be genuine; novel transcripts were then assembled with StringTie [53], retaining only transcripts representing more than 5% of expression from a locus (-f 0.05). Assemblies from individual samples were then merged, keeping only transcripts with a TPM of greater than 1.

Once this pipeline had been run on each cancer type individually, all assemblies were merged into a master “all_TCGA” assembly, to facilitate comparison between different cancer types downstream. 3UIs are then detected by comparison against a high confidence reference assembly (Ensembl85; transcript support level 1 or 2 and having an APRIS primary or alternate 1 or 2 annotation) to classify 3UIs. Briefly, each transcript in the assembly is compared to each transcript in the reference set that has a start codon in an exon of the query transcript. A 3UI transcript is identified when the transcript shares identical intron chains within the boundaries of the reference transcripts CDS, and additional introns 3’ of the reference stop codon. Transcripts are classed as ‘nonPTC’ if the 3UI does not share an exon-intron or intron-exon boundary with a coding intron in any reference transcript.

Transcripts are classified as ‘novel’ if they contain a 3UI that does not share any exon-intron or intron-exon junction with any reference transcript. The distance of the 3UI from the stop codon is recorded as the distance from the stop codon of the matched reference transcript. Genesets were compared using trmap (https://github.com/gpertea/trmap). This pipeline can be found at www.github.com/sudlab/Cancer3UIs/pipelines/pipeline_utrons_annotate.py.

We checked for spurious assembly of 3UI transcripts by simulating 60 samples of RNAseq reads from the Ensembl 85 annotation using Polyester [54], with all transcripts simulated at 1TPM, and passing the resulting files through the pipeline outlined above. Any gene for which a transcript with a novel 3UI was identified by this process was excluded from further analyses.

Once pan-TCGA and pan-CCLE assemblies were obtained, samples were then quantified against the appropriate assembly using Salmon (with options --gcBias --reduceGCMemory) with an index using the full sequence of hg38 as decoy, and reads mapping to exons and junctions counted using featureCounts [55] (with options -fJOG). Transcripts were classed as “broadly expressed” in a given tissue where they had a Transcripts Per Million (TPM) value > 1 and fraction expression (Transcript TPM ÷ Gene TPM) > 0.25 in at least 10% of either cancer or normal samples, and evidence of all junctions being covered by at least one read from featureCounts. This pipeline can be found at www.github.com/sudlab/Cancer3UIs/pipelines/pipeline_utrons_requant.py.

### Correlation of average 3UI PSO vs gene expression per sample

For this analysis individual event level PSO (percent spliced out; 1-PSI) values for normal and cancerous colon tissue from TCGA were summarised to produce average 3UI PSO values for both nonPTC 3UIs and pPTC 3UIs, per sample. The average 3UI PSO values were then correlated with the normalised gene expression value of every protein coding gene in the genome. Normalisation of gene expression was conducted using the DESeq2 method [56]. To test the strength and significance of correlations between normalised gene expression and average 3UI PSO, Spearman’s rank correlation coefficient was calculated per gene per condition (normal nonPTC, normal pPTC, cancer nonPTC, cancer pPTC). P-values were adjusted via Benjamini-Hochberg correction.

### Event- and transcript-level splicing analyses

For Differential Transcript Usage (DTU) analysis DRIMSeq [57] was used. Transcripts were only considered for DTU analyses where they had counts > 10 and fraction expression > 0.1, in > 10% of samples. The output of DRIMSeq was filtered for 3UI-containing transcripts before p-value adjustment, adjusted p-values ≤ 0.05 were considered statistically significant.

rMATS-turbo [58] was used for differential splicing analysis against a retained intron fixed event set generated from our ‘all_TCGA’ transcript assembly. Our rMATS pipeline can be found at www.github.com/sudlab/Cancer3UIs. For analysis of TCGA RNAseq data each cancer type was analysed separately. Inputs were provided in .bam format, where ‘b1’ was set as cancer samples and ‘b2’ was set as normal samples, meaning events with IncLevelDifference < 0 are more spliced in cancer than in normal. For Wnt manipulation RNAseq data we set ‘b1’ as 0μM CHIR99021 and ‘b2’ as 20μM CHIR99021, therefore events with IncLevelDifference > 0 are more spliced in 20μM CHIR99021 treated. Only results produced from junction counts (not exon counts) were taken for downstream analysis. Events were considered significant where FDR<0.1. Sashimi plots were generated using ‘rmats2sashimiplot’ (www.github.com/Xinglab/rmats2sashimiplot).

### RNA and miRNA enrichment analyses

RNA binding protein and miRNA interaction annotations were downloaded from starBase/ENCORI [59]. These were subsequently compared with a .bed file containing the locations of nonPTC 3UIs using ‘GAT’ [60]. Fold change is a representation of observed motif frequency compared with expected frequency (which is generated through randomized sampling). Enrichment was considered statistically significant where adjusted p-values ≤ 0.01.

### Cell culture and treatments

Human colorectal carcinoma cell line HCT116 was cultured in High Glucose DMEM (4.5g/L) supplemented with 10% fetal calf serum and Penicillin-Streptomycin and maintained at 37°C and 5% CO_2_ (standard conditions). For Wnt manipulation assays HCT116 were initially grown for 24 hours under standard conditions before media was changed to RPMI 1640 supplemented with varying concentrations (0-20μM) of CHIR99021 (for 24 hours) or IWR-1 (for 48 hours).

### RNA extraction, gDNA extraction, RT-PCR, RT-qPCR, RNAseq

RNA extraction was conducted using TRI Reagent (Sigma-Aldrich) in line with the manufacturer’s instructions. DNA removal was facilitated by TURBO DNAse (Invitrogen) treatment for 1 hour, in the presence of RiboSafe RNase Inhibitor (Bioline). Reverse transcription was conducted using High-Capacity cDNA Reverse Transcription Kit (Thermo Fisher) in the presence of RiboSafe RNase Inhibitor. gDNA was extracted by lysing and incubating cells in 199μl 1M Tris-EDTA, 0.5μl 20% sodium dodecyl sulfate, and 0.52μl 19mg/ml proteinase K for 4 hours at 60°C with shaking, followed by phenol-chloroform extraction and EtOH precipitation.

RT-PCR was conducted using Quick-Load Taq 2X Master Mix (NEB). For molecular cloning applications Q5 High-Fidelity DNA polymerase (NEB) was used. RT-qPCR was conducted using SensiMix SYBR Hi-ROX 2x Master Mix (Bioline) on the Rotor-Gene Q (Qiagen). For sequencing applications RNA was extracted as previously described, quality was assessed by Qubit RNA Broad-Range Assay Kit (Thermo Fisher), mRNA library preparation and Illumina short-read (PE150) sequencing was conducted by Novogene (Cambridge, UK).

### NMD inhibition

HCT116 cells were seeded 24 hours prior to siRNA transfection to achieve 30% confluency at the time of transfection. siRNA duplexes against UPF1 (UPF1_1: CAGUUCCGCUCCAU UUUGAU; UPF1_2: GAUGCAGUUCCGCUCCAUU) were transfected at 30nM using Lipofectamine RNAiMAX (Invitrogen) as per the manufacturer’s instructions. Cells were transfected in the same manner 48 hours later and harvested 72 hours following initial transfection. Efficiency of UPF1 knockdown was assessed by Western blotting and RT-qPCR. For small molecule NMD inhibition, UPF1 inhibitor NMDI14 (Sigma-Aldrich #SML1538) was supplemented into culture media 24 hours prior to RNA extraction at a final concentration of 65μM.

### Western blotting

20µg protein was resolved by SDS-PAGE in a Mini-PROTEAN vertical electrophoresis cell (Bio-Rad) alongside PageRuler Prestained Protein Ladder (Thermo Scientific #26619). Protein transfer was performed onto nitrocellulose membranes using the Trans-Blot Turbo (Bio-Rad) at 25V, 1.3mA for 25 minutes. Membranes were blocked with 5% milk in TBST for 1 hour at room temperature followed by incubation with 1:500 anti-UPF1 antibody (Proteintech #66898) or 1:5000 anti-Tubulin (Sigma-Aldrich #T6199) in TBST + 5% milk for 2 hours at room temperature. Membranes were washed 3X with TBST for 5 minutes. Membranes were then incubated with 1:10000 HRP-conjugated secondary antibody (Promega #W402B) in TBST + 5% milk for 1 hour at room temperature. Membranes were washed 3X with TBST for 5 minutes. Signal was developed by ECL reagent for 30 seconds before the membrane was exposed using a BioRad Chemidoc.

### Molecular cloning

Full length CTNNB1 and HRAS 3’ UTRs were amplified using Q5 High-Fidelity DNA polymerase from HCT116 gDNA. Following DpnI digest and gel extraction, PCR products were cloned into pCI-neo (Promega) downstream of the Luciferase2 gene under the control of a CMV promoter. Q5 Site-directed mutagenesis kit (#E0554S, NEB) was used to produce intronless, 5’ss and 3’ss mutants as per the manufacturer’s instructions. Plasmids were verified by Sanger sequencing.

### Luciferase assays

HCT116 cells were seeded at 125000 cells per well into 24-well plates. Cells were transfected after 24 hours with 500ng of Luc2-CTNNB1/HRAS and 5ng hRluc plasmids using polyethylenimine (PEI) at a 1:3 DNA:PEI ratio. Transfected cells were cultured under standard conditions for 72 hours before being washed with PBS and lysed and processed using materials provided with the Dual-Luciferase Reporter Assay System Kit (Promega).

### RNA stability assays

HCT116 cells were seeded at 300000 cells per well into 6-well plates. After 48 hours culture media was supplemented with 5ug/ml actinomycin D. Cells were harvested at 0hrs, 3hrs, 12hrs following the addition of actinomycin D. RNA was extracted with Total RNA Purification Plus Kit (#48400; Norgen Biotek). Isoform expression was measured via qPCR following cDNA synthesis (previously described).

### Differential transcript expression analysis

Differential transcript expression analysis for UPF1 knockdown in HCT116, CHIR99021 treatment in HCT116, and nucleocytoplasmic fractionation in HCT116 [61] (GSE228810) were conducted using DESeq2. Principal component analysis (PCA) was performed as a method of quality control. For UPF1 knockdown in HCT116 PCA indicated no effect of the small molecule inhibitor, therefore the primary coefficient used in this analysis was siUPF1 vs siDsRed. ECDF (empirical cumulative distribution function) plots were generated to compare Log2FoldChange trends between subsets of transcripts. For nucleocytoplasmic fractionation data analysis a prior filtering step was conducted to ensure that only transcripts from protein coding genes were analysed, and that these had a transcript expression of >1 TPM in either the nuclear or cytoplasmic fraction.

## Supporting information

Additional File 4 - non-PTC CCLE 3UIs

Additional File 5 - All TCGA 3UIs

Additional File 6 - non-PTC TCGA 3UIs

Additional File 7 - TCGA transcript assembly

Additional File 9 - Broadly Expressed Transcripts

Additional File 1 - All CHESS 3UIs

Additional File 2 - non-PTC CHESS 3UIs

Additional File 3 - All CCLE 3UIs

## Abbreviations

3’ UTR: 3’ Untranslated region
3UI: 3’ UTR intron
APA: Alternative polyadenylation
AS: Alternative splicing
CDS: Coding sequence
DEU: Differential exon usage
DTU: Differential transcript usage
EJC: Exon junction complex
MRE: miRNA recognition element
NMD: Nonsense-mediated decay
PTC: Premature termination codon
RBP: RNA binding protein
TPM: Transcript per million

## Declarations

### Ethics approval and consent to participate

Not applicable

### Consent for publication

Not applicable

### Availability of data and materials

Raw sequencing datasets and corresponding salmon quantification files have been submitted to the NCBI Gene Expression Omnibus (GEO) under accession numbers GSE251665 and GSE251666. Transcript and gene quantifications, fraction expression calculations, exon counts, junction counts, PSIs and differential splicing data for TCGA samples is available from Sheffield Online Research Data Archive (ORDA) under DOI 10.15131/shef.data.25612722. Custom pipelines and scripts (alongside software versions) used herein, are available at: https://github.com/sudlab/Cancer3UIs.

### Authors’ contributions

The study was conceived by IS and SW. IS, JR, CA-C and SB-S designed and carried out bioinformatic analysis under the supervision of IS. UPF1 knockout RNA-seq experiment was designed and carried out by CA-C under the supervision of SW. All other experiments were designed by IS, JR and SW, and carried out by JR under the supervision of SW. IS and JR wrote the manuscript with input from all authors.

## Acknowledgements

The results published here are in whole or part based upon data generated by the TCGA Research Network: https://www.cancer.gov/tcga. We acknowledge IT Services at The University of Sheffield for the provision of services for High Performance Computing.

