## Additional File 8 - Supplementary Figures for "Widespread 3’ UTR splicing regulates expression of oncogene transcripts in sequence-dependent and independent manners"

**Supplementary Figure 1. 3'UTR splice site conservation and examples.** (A) Percentage of 3'UTR splice sites that utilize the canonical GT-AG splice site versus non-canonical sites. (B) Conservation of 5' and 3' splice site sequence compared to surrounding sequence. (C) RT-PCR using primers flanking the 3'UTR intron for HRAS, SRSF8 and CTNNB1, amplified from either HCT116 cDNA or gDNA.

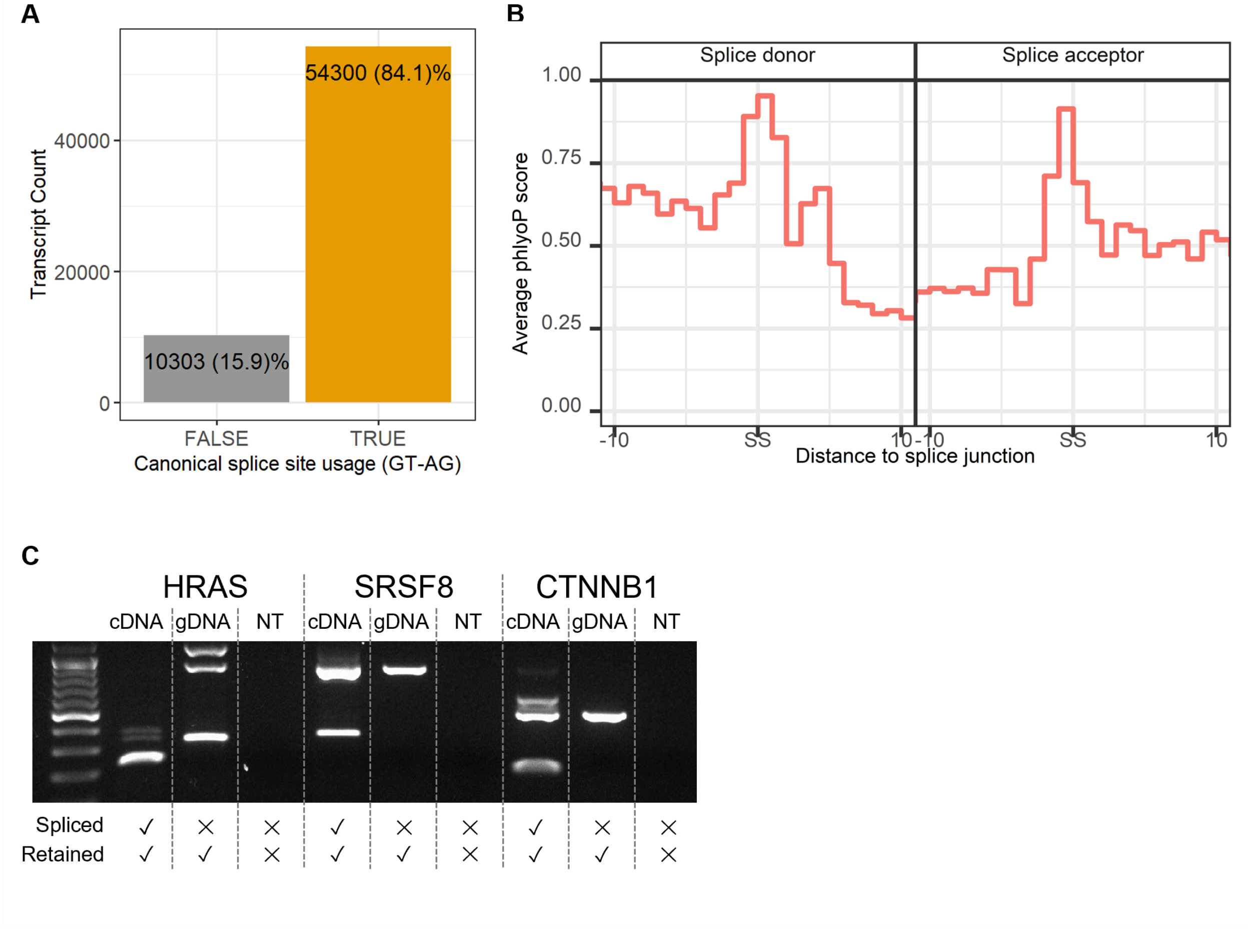

**Supplementary Figure 2. Saturation analysis and artefact simulation.** (A) Saturation analysis of 3UI detection utilising an increasing number of randomly sampled colon cancer RNA sequencing samples from TCGA shows detection begins to saturate at 100 samples. B) Number of novel 3UIs identified in 60 simulated RNA-seq datasets. Plot shows the number of transcripts and number of genes containing novel 3UIs for successively larger numbers of simulations

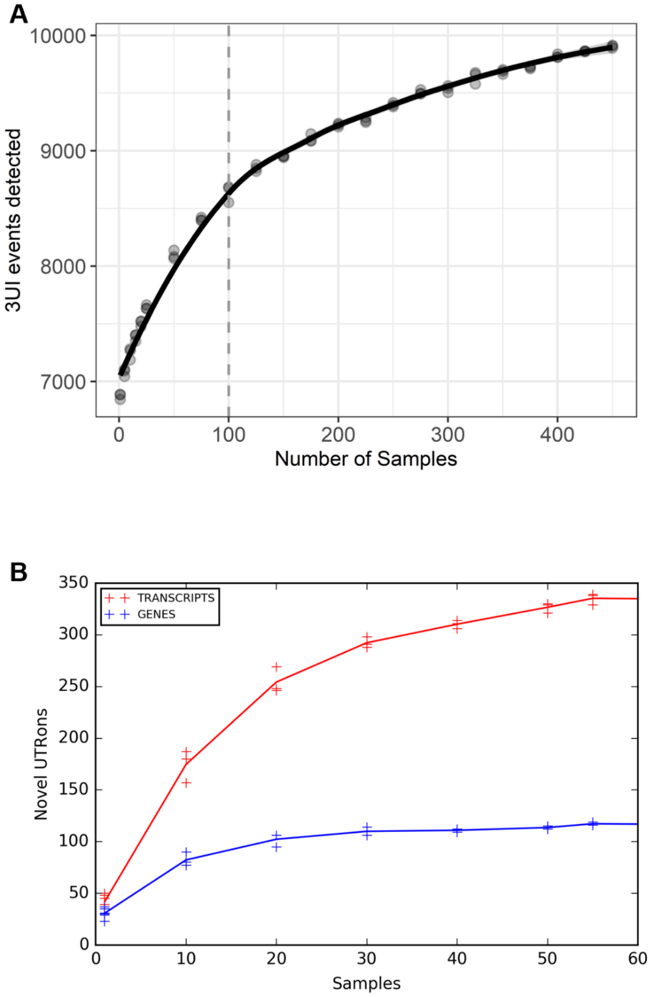

**Supplementary Figure 3. Broad expression of 3UI-containing transcripts.** Curve to represent the number of transcripts that meet each expression criteria in X% of colon cancer samples. Each colour represents an expression cutoff.

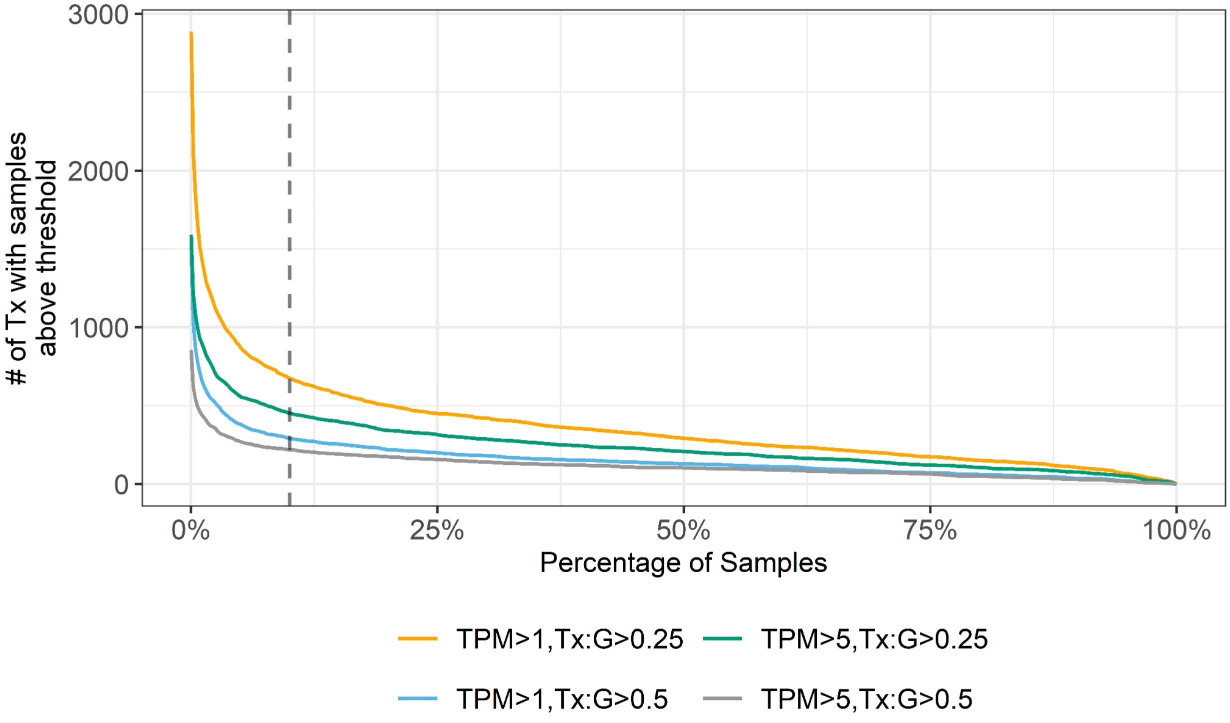

**Supplementary Figure 4. Average 3UI PSO correlations.** (A) Heatmap displaying the differences in correlation coefficient estimates between normal vs cancer and nPTC vs pPTC samples for NMD components. (B) Correlation of average 3UI PSO with normalised UPF1 expression. (C) Correlation of average 3UI PSO with normalised UPF3B expression. (D) Correlation of average 3UI PSO with normalised SMG8 expression.

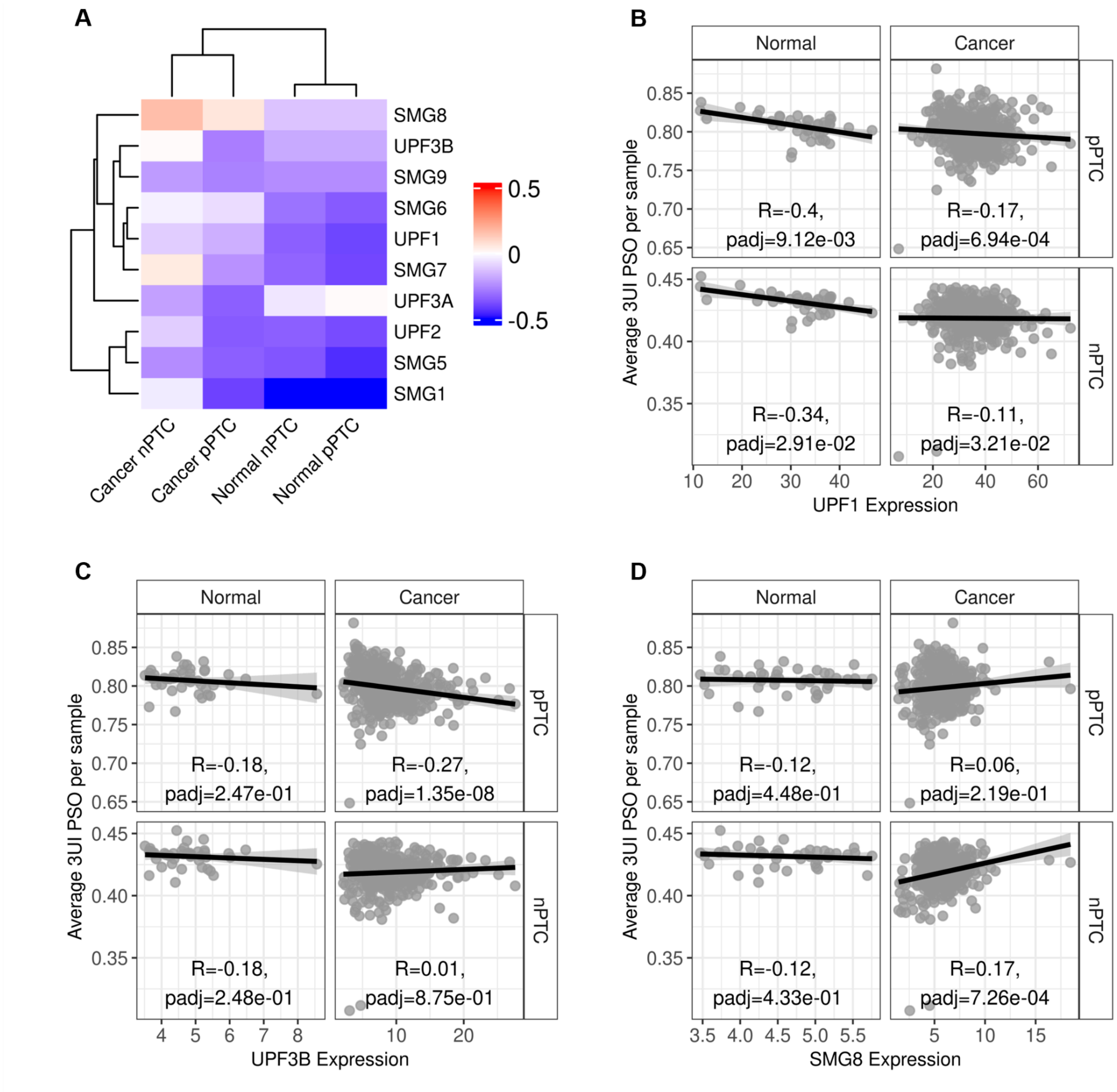

**Supplementary Figure 5. UPF1 knockdown validation and effect on HRAS isoforms.** (A) Western blot against UPF1 and Tubulin for HCT116 cells transfected with either siUPF1_1, siUPF1_2 or siDsRed (control). (B) RT-qPCR validation of UPF1 knockdown. (C) HRAS retained and spliced isoform expression measured by RT-qPCR following transfection with either siUPF1_2 or siDsRed (control). (D) HRAS retained and spliced isoform expression from RNAseq data following transfection with either siUPF1_1 or siDsRed (control).

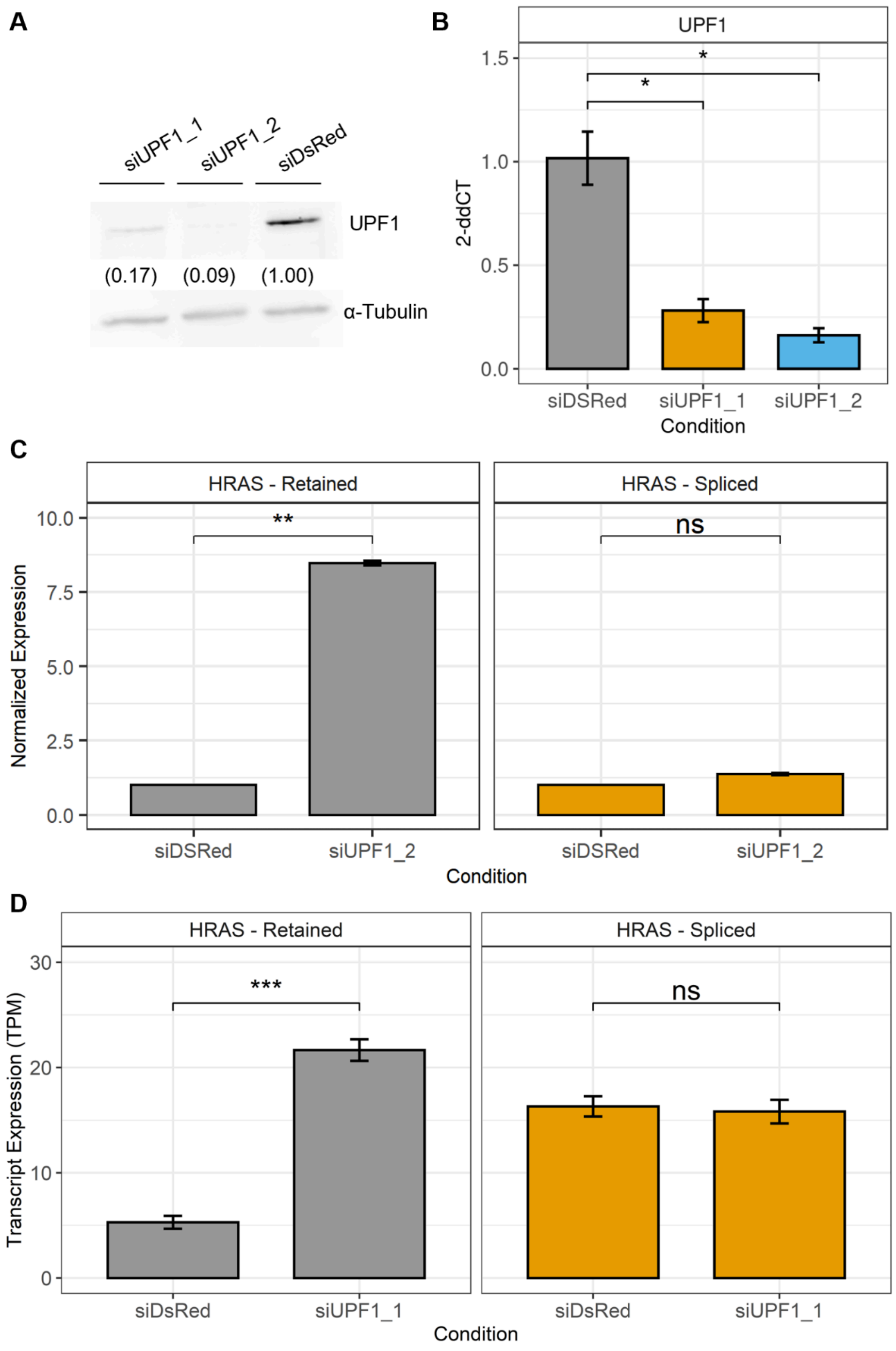

**Supplementary Figure 6. Luciferase 3'UTR reporter plasmids.** (A) Splicing of Luciferase2-HRAS 3'UTR plasmids upon transfection into HCT116 cells. FL produces both retained (R) and spliced (S) isoforms. ΔI produces only S. 5'ss mutant produces only R. (B) Schematic representation of Luciferase2-CTNNB1 3'UTR plasmids. (C) Splicing of Luciferase2-CTNNB1 plasmids upon transfection into HCT116 cells. FL produces both retained (R) and short-spliced (S) isoform, long-spliced (L) is not visible. 5'ss mutant produces R only. ΔI(S) produces S only. ΔI(L) produces L only. 3'ss(S) produces R and L. 3'ss(L) produces R and S.

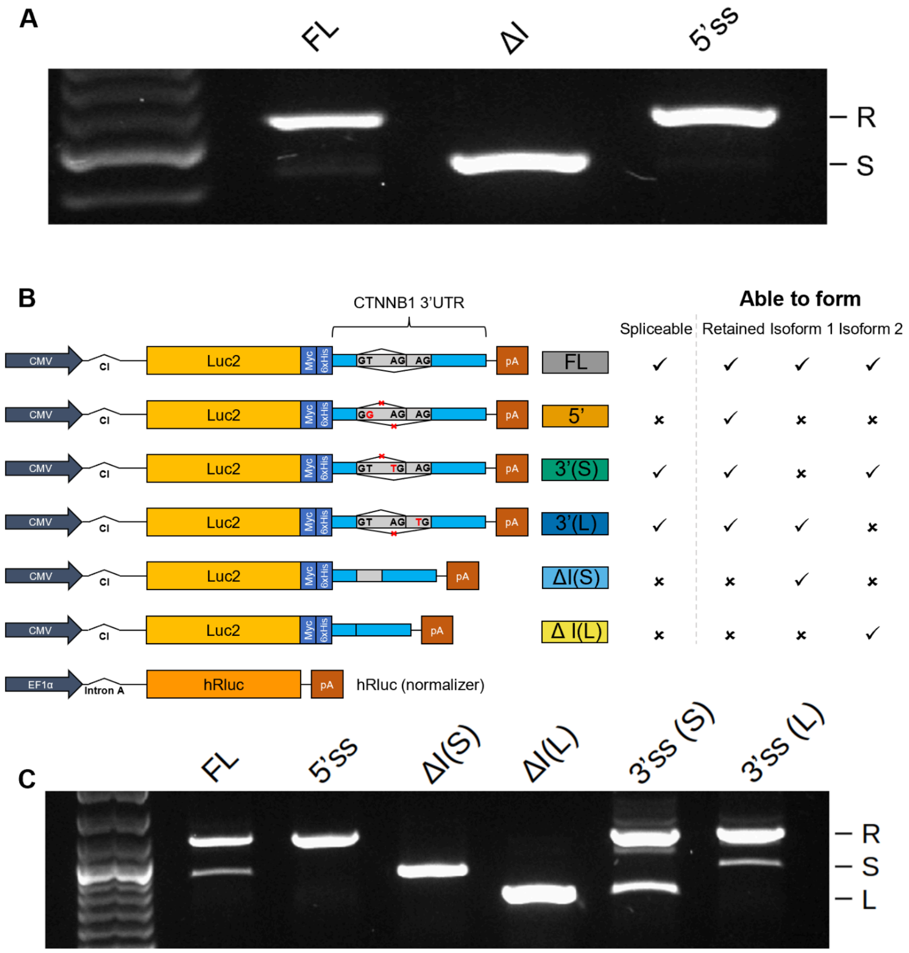

**Supplementary Figure 7. Differential 3'UTR splicing between normal and cancer samples.** Distribution of IncLevelDifferences for 3UIs between normal and cancer samples. Grey density plots represent nonPTC 3UIs. Orange density plots represent pPTC 3UIs.

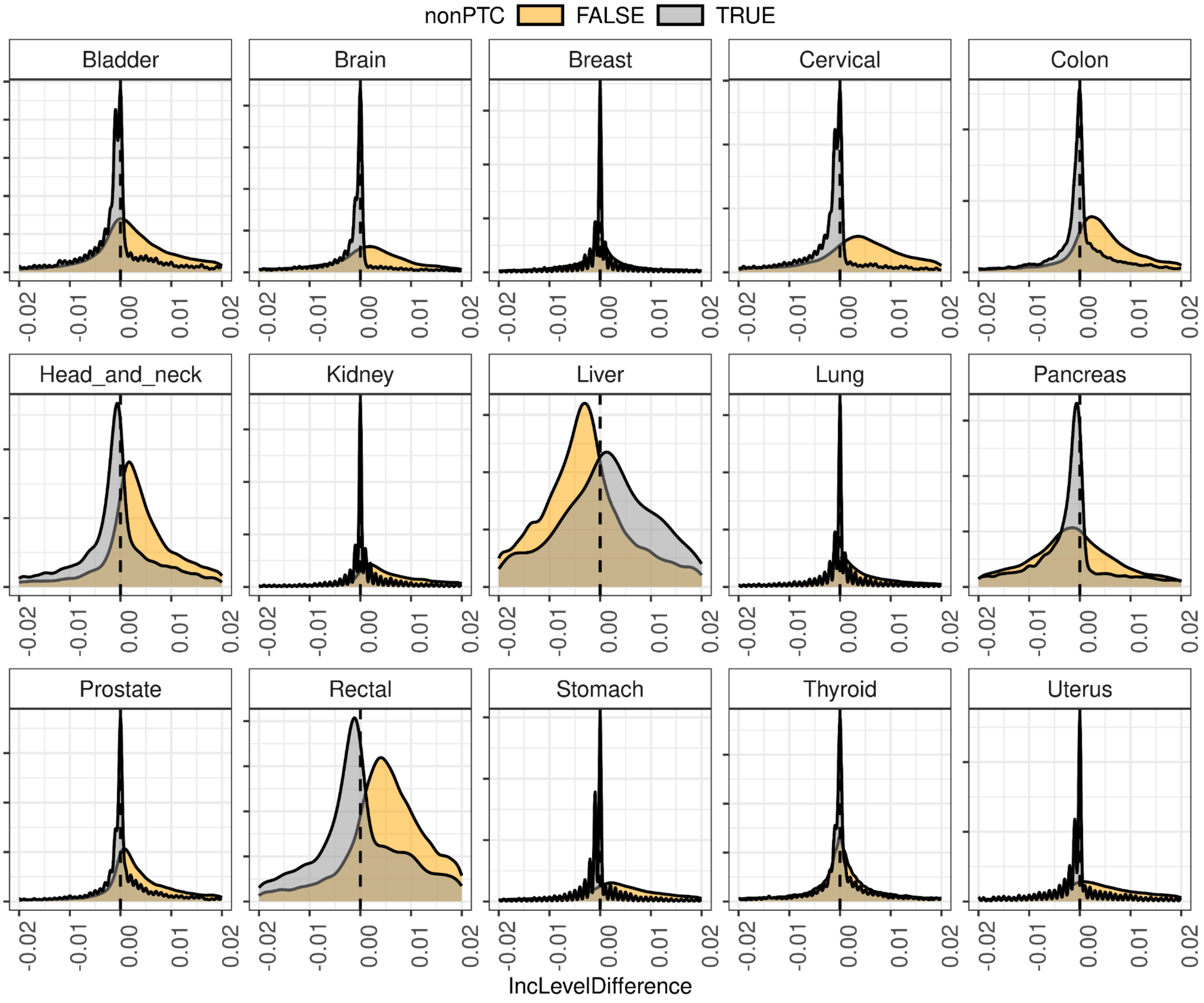

**Supplementary Figure 8. RBP and miRNA enrichment analysis.** (A) Enrichment of RBPs in all detected nonPTC 3UIs, supported by RBP-CLIPseq data. (B) Enrichment of miRNAs in all detected nonPTC 3UIs, supported by AGO-CLIP data.

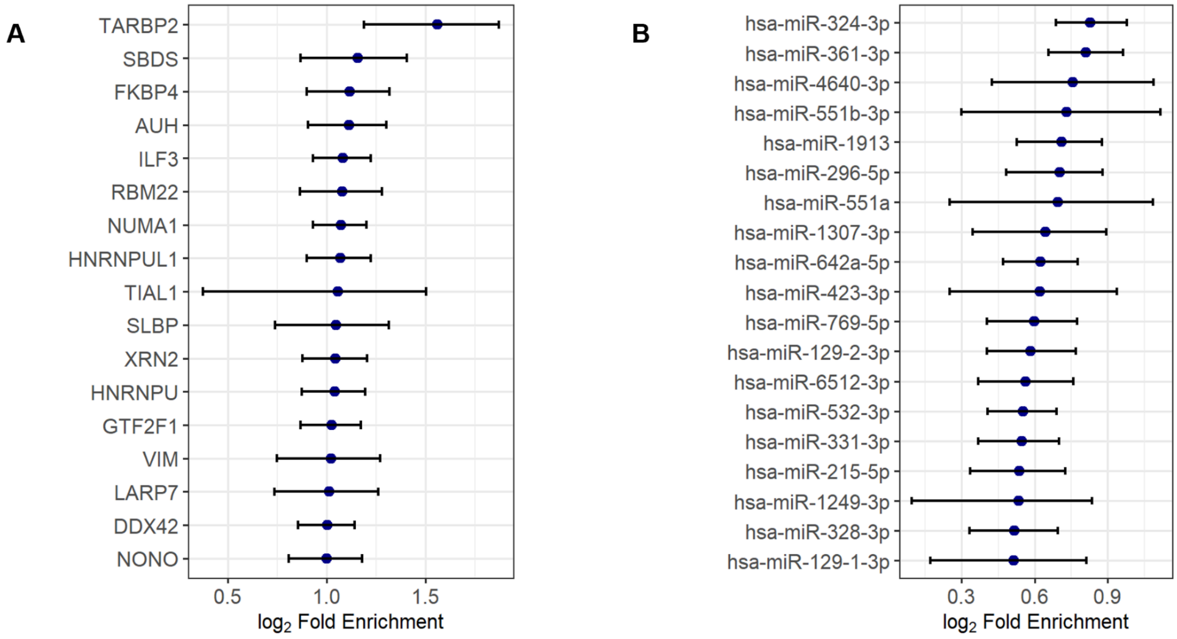

**Supplementary Figure 9. 3’UTR splicing of the Canonical Wnt signalling pathway.** Breakdown of the canonical Wnt signalling pathway colour-coded by whether each component 3’UTR is spliced more (orange) or retained more (blue) upon Wnt signalling activation in HCT116 cells. P-value derived from Gene Set Enrichment Analysis.

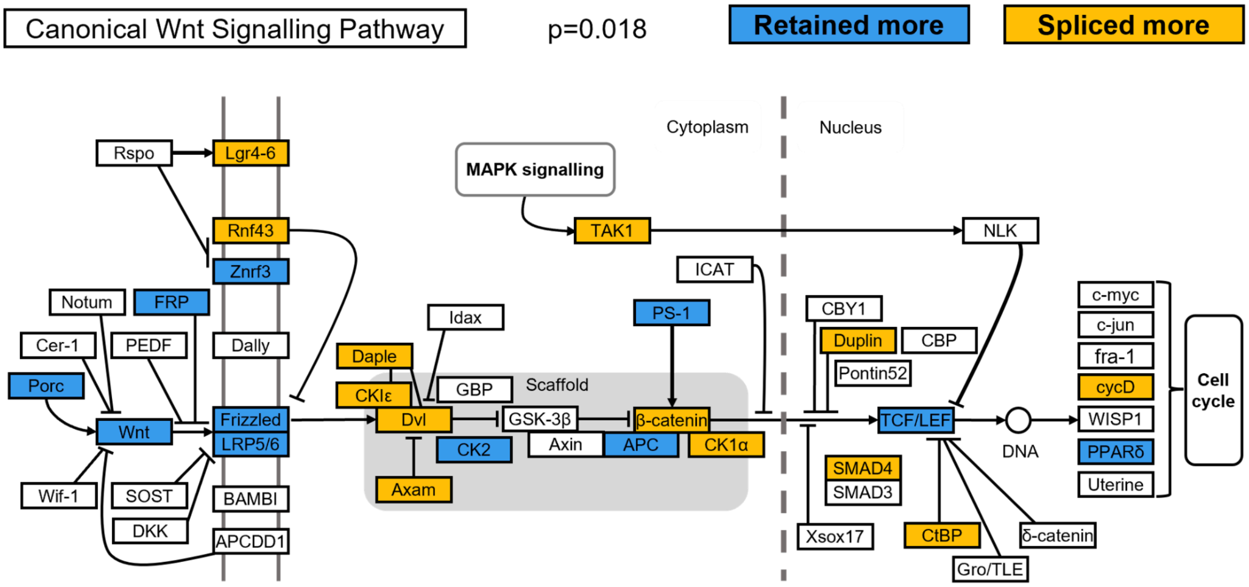

**Supplementary Figure 10. Schematic model explaining Luciferase assay results.** (A) Where the intron is cloned out via molecular cloning as opposed to splicing endogenously, no EJC will be deposited, therefore endogenous EJC-dependent regulatory pathways will impact full-length but not ΔI constructs. These may increase or decrease stability depending on the mRNP composition. (B) Through cloning out an intron we may introduce a cis-element which can bind trans-factors such as miRNAs or RBPs, which would otherwise be non-functional in the endogenous context due to the presence of the EJC.

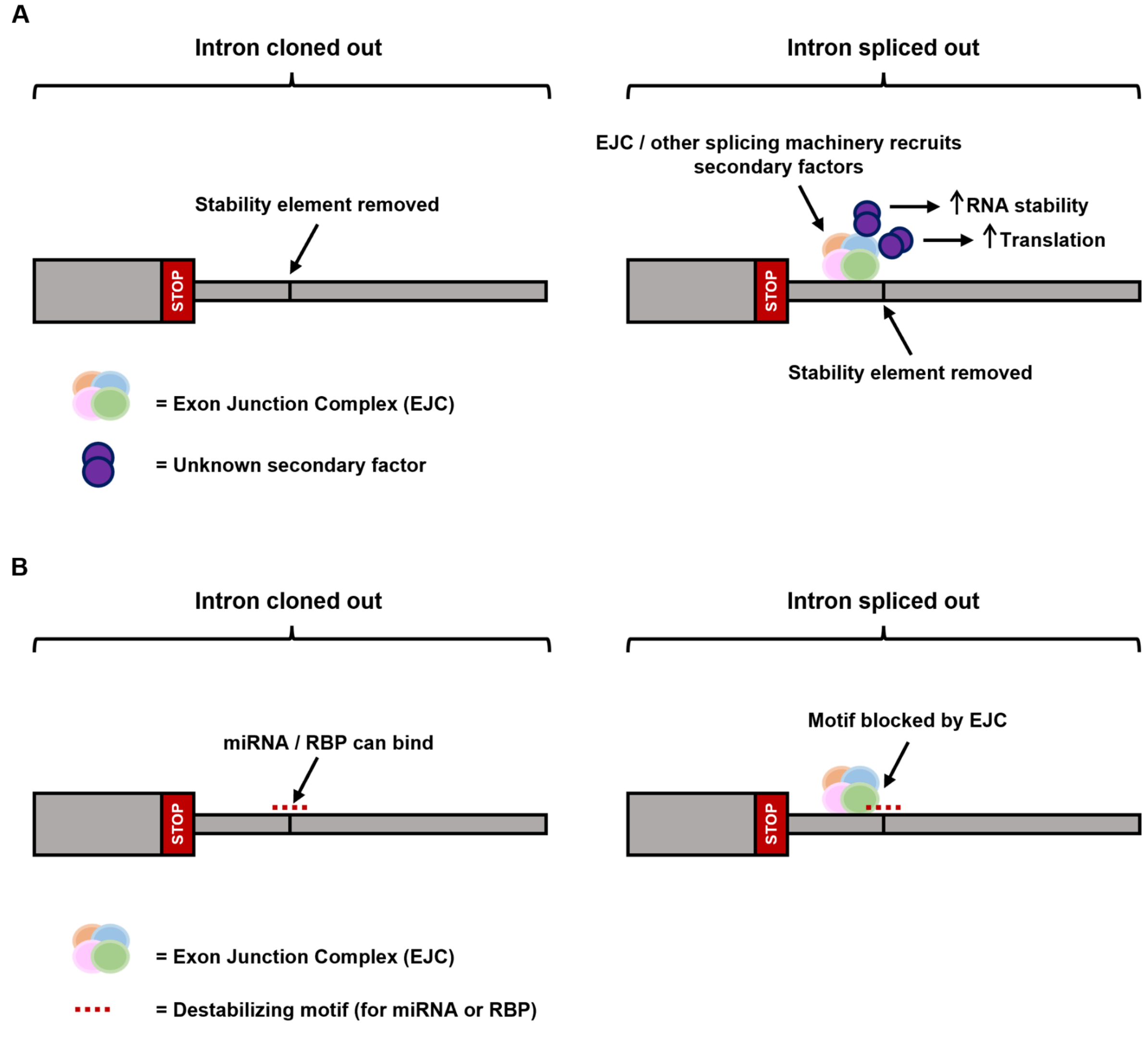

**Supplementary Figure 11. Transcript stabilisation by 3'UTR splicing.** Splicing 3'UTRs may remove sequences that would otherwise be subject to m6A modification, which is predominantly a destabilising marker. Additionally, the presence of the EJC upstream of the splice site prevents m6A deposition in the local vicinity, as part of a so-called "m6A exclusion zone".

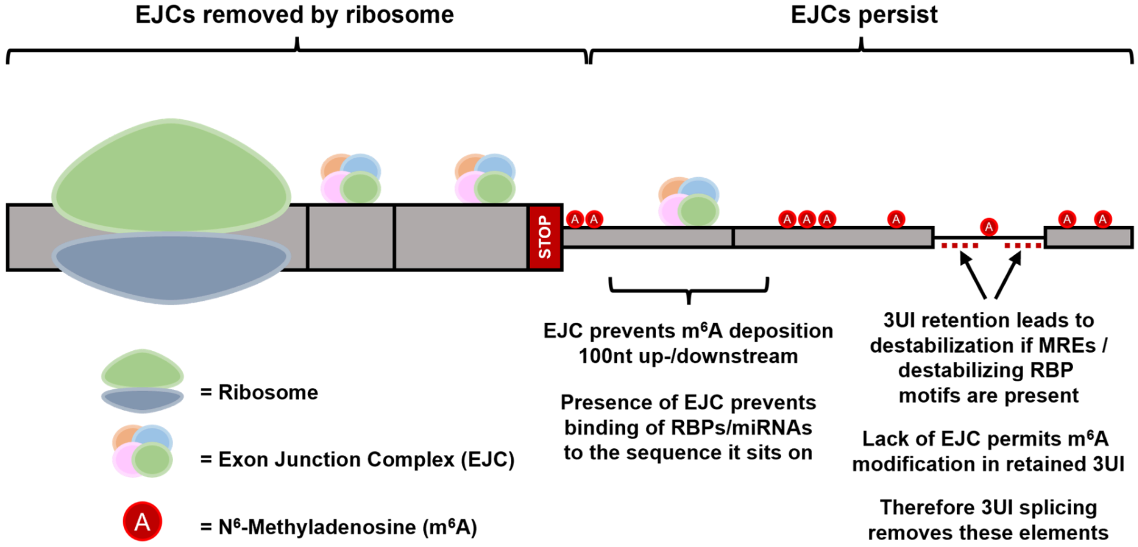
